# The thermodynamics of thinking: connections between neural activity, energy metabolism and blood flow

**DOI:** 10.1101/833855

**Authors:** Richard B. Buxton

**Affiliations:** University of California San Diego

**Keywords:** functional neuroimaging, cerebral blood flow (CBF), cerebral metabolic rate of oxygen (CMRO2), neural activity, thermodynamic limitations, hypoxia

## Abstract

Several current functional neuroimaging methods are sensitive to cerebral metabolism and cerebral blood flow (CBF) rather than the underlying neural activity itself. Empirically, the connections between metabolism, flow and neural activity are complex and somewhat counterintuitive: CBF and glycolysis increase more than seems to be needed to provide oxygen and pyruvate for oxidative metabolism, and the oxygen extraction fraction is relatively low in the brain and *decreases* when oxygen metabolism increases. This work lays a foundation for the idea that this unexpected pattern of physiological changes is consistent with basic thermodynamic considerations related to metabolism. In the context of this thermodynamic framework, the apparent mismatches in metabolic rates and CBF are related to preserving the entropy change of oxidative metabolism, specifically the O_2_/CO_2_ ratio in the mitochondria. However, the mechanism supporting this CBF response is likely not due to feedback from a hypothetical O_2_ sensor in tissue, but rather is consistent with feed-forward control by signals from both excitatory and inhibitory neural activity. Quantitative predictions of the thermodynamic framework, based on models of O_2_ and CO_2_ transport and possible neural drivers of CBF control, are in good agreement with a wide range of experimental data, including responses to neural activation, hypercapnia, hypoxia and high-altitude acclimatization.

## 1. Introduction: The challenge of interpreting metabolism and blood flow dynamics in terms of the underlying neural activity

Current functional non-invasive neuroimaging methods such as functional magnetic resonance imaging (fMRI), positron emission tomography (PET), and near infrared spectroscopy (NIRS) do not measure neural activity directly, but instead are sensitive to metabolic and blood flow changes that accompany changes in neural activity [1]. Consequently, understanding the links between metabolism, flow and neural activity is an active goal of current neuroimaging research. Intuitively, we expect strong connections because neural activity is energetically costly [2-10]. To simplify a complex process, excitatory synaptic activity generates sodium currents from the extracellular to intracellular space, and in recovery from this activity neurons must then pump sodium back out of the cell against a steep thermodynamic gradient. This is accomplished by coupling sodium transport to the conversion of adenosine triphosphate (ATP) to adenosine diphosphate (ADP), a thermodynamically highly favorable reaction, within the sodium/potassium pump [11]. ATP, the ubiquitous energy currency of the cell, is then restored through glucose metabolism [8, 12, 13]. The first step is glycolysis in the cytosol, generating a small amount of ATP with the conversion of glucose to pyruvate (Pyr). Much more ATP is then generated by oxidative metabolism of Pyr in the mitochondria, consuming O_2_ and producing CO_2_. Blood flow delivers the O_2_ and glucose and carries away the CO_2_.

The simplest scenario we might have imagined for this net process would be a serial chain of events with each triggering the next: recovery from neural activity depletes ATP, reduced ATP stimulates glycolysis and oxidative metabolism, and increased metabolism stimulates blood flow. In addition to this simple chain of driving mechanisms, we might have anticipated proportional changes of the cerebral metabolic rate of glucose (CMRGlc), the cerebral metabolic rate of oxygen (CMRO_2_), and cerebral blood flow (CBF), all matched to the degree of neural activity change. Instead, though, current research suggests a much more complicated picture (**Figure 1, upper panel**), with several counterintuitive features, and it is the complexity of this process that currently presents a barrier to more quantitative interpretations of metabolism and flow dynamics in terms of the underlying neural dynamics.

**Figure 1.**
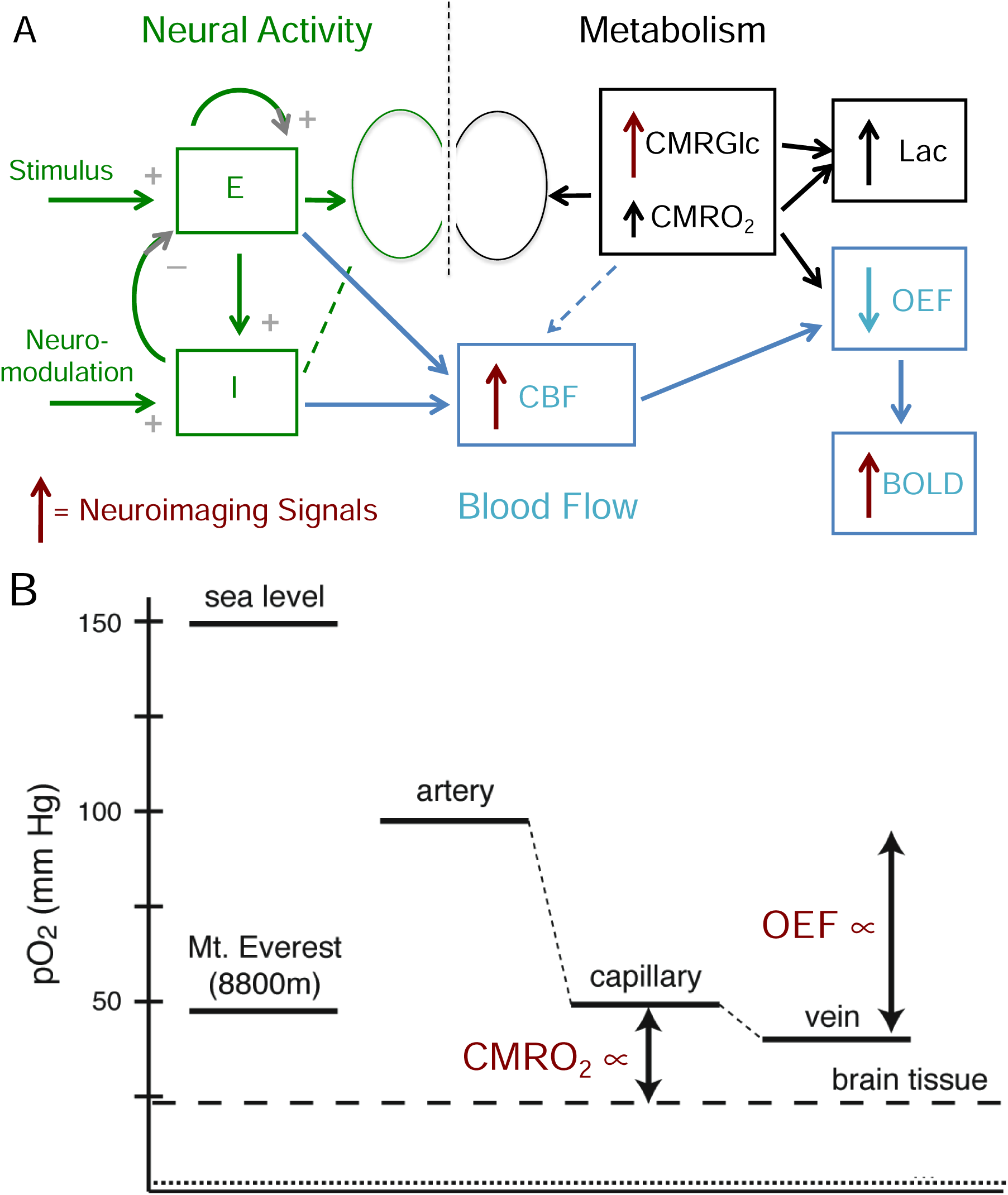
Schematic of the connections between neural activity, metabolism, blood flow and neuroimaging signals. **Upper Panel:** Input stimulus and neuromodulatory signals evoke activity of interacting excitatory (E) and inhibitory (I) neural populations (left, green), and ATP is consumed in recovery from that activity, primarily in restoring ion gradients. The ATP is restored through oxidative metabolism of glucose (right, black), with oxygen and glucose delivered by blood flow (middle, blue; CMRGlc = cerebral metabolic rate of glucose (glycolysis), CMRO_2_ = cerebral metabolic rate of O_2_, CBF = cerebral blood flow). Solid/dashed connecting arrows reflect strong/weak drivers, respectively, and the vertical arrows reflect the fractional increase of each rate. The larger increase of glycolysis than CMRO_2_ leads to increased lactate production (Lac), and the larger increase of CBF than CMRO_2_ leads to decreased oxygen extraction fraction (OEF), which produces the blood oxygenation level dependent (BOLD) signal measured in fMRI. The sources for current neuroimaging signals (fMRI and PET) are indicated by red arrows. The seemingly paradoxical features addressed in this paper are: the mismatch of CMRGlc and CMRO_2_, leading to increased Lac production despite the availability of O_2_; the mismatch of CBF and CMRO_2_, leading to the decreased OEF; and the presence of a strong feed-forward drive from the inhibitory neural population (I) to CBF, even though the I activity is likely to be only a small contribution to the ATP costs and thus CMRO_2_. **Lower Panel:** Diagram of O_2_ partial pressures, with atmospheric values on the left for comparison and physiological values on the right. To increase CMRO_2_, the diffusion gradient of PO_2_ between the capillary blood and the mitochondria (tissue) must increase. A consequence of increasing CBF more than CMRO_2_ is that the oxygen extraction fraction (OEF) is reduced, raising capillary PO_2_. In this way the PO_2_ gradient can be increased, supporting increased CMRO_2_, while maintaining the tissue PO_2_ at a constant level. The larger increase of CMRGlc than CMRO_2_ may serve an analogous function by increasing cytosolic Pyr to increase the diffusion gradient to the mitochondria without reducing mitochondrial Pyr. In the context of the proposed thermodynamic framework, the large increases of CBF and CMRGlc serve the broader role of maintaining the entropy change of oxidative metabolism. For CBF, a key element is that it controls the tissue O_2_/CO_2_ level, and this has additional implications for the CBF response to hypercapnia and hypoxia.

The most striking departure from the simple picture described above is the apparent mismatch of the rates of different processes. The CMRGlc change with increasing neural activity is much larger than the CMRO_2_ change [8, 13-15], and some of the excess pyruvate created is converted to lactate [15, 16]. The production of lactate is surprising given that there appears to be no lack of available oxygen, and this phenomenon has been called ‘aerobic glycolysis’ [8, 13]. From the perspective of increasing ATP production, the added glycolysis has little impact because the ATP production from oxidative metabolism in the mitochondria is more than 15 times the ATP production from glycolysis alone. For example, even a 50% increase of glycolysis alone would increase ATP production by only about 3%. In short, even with this added glycolysis most of the needed increase in ATP production is due to the increased oxidative metabolism, reflected in CMRO_2_ [17, 18].

Blood flow also increases much more than CMRO_2_ [19-32], so that oxygen delivery to tissue is increased much more than the actual increase in the rate at which it is metabolized. A natural idea is that the large blood flow increase might be necessary to support the large change in CMRGlc, even though the function served by the large change in glycolysis is still unclear. However, studies limiting the increase of CBF nevertheless found normal increases of CMRGlc [33, 34], indicating that the large change in CBF is not necessary to support CMRGlc.

The phenomenon of a much larger increase of CBF than CMRO_2_ creates the seemingly paradoxical effect that the oxygen extraction fraction (OEF), the fraction of delivered oxygen that is extracted and metabolized, *decreases* with increased neural activity, and this effect is at the heart of the blood oxygenation level dependent (BOLD) changes of the measured fMRI signal [35, 36]. The physical origin of the BOLD effect is that deoxyhemoglobin has paramagnetic properties that create magnetic field distortions and reduce the measured MR signal [37]. The reduction of OEF with increased neural activity then reduces those field distortions, creating a slight increase of the MR signal. Importantly, the magnitude of the BOLD signal depends on the degree of mismatch between CBF and CMRO_2_, as well as the amount of deoxyhemoglobin present in the baseline state, creating a critical challenge to any quantitative interpretation of the BOLD signal alone in neural terms [37, 38]. In short, the mismatch of CBF and CMRO_2_ makes possible BOLD-fMRI, but what function is served by this mismatch, and under what circumstances might the degree of mismatch change?

The primary idea for understanding this mismatch is that a larger CBF increase than the CMRO_2_ increase tends to maintain the tissue O_2_ level [39-42]. As CMRO_2_ increases, the gradient of O_2_ concentration from blood to tissue must increase to support a higher diffusive flux of O_2_ to the mitochondria (**Figure 1, lower panel**). In principle, this gradient could be increased by lowering the partial pressure of oxygen (PO_2_) in tissue or by raising blood PO_2_, and we made the early suggestion that tissue O_2_ concentration is so low to begin with that the only option is to raise blood O_2_ [43]. By reducing the OEF, the capillary PO_2_ is increased, increasing the PO_2_ gradient from blood to mitochondria. Later studies further developed this idea in the context of non-zero tissue PO_2_ [44-48], with a less severe requirement for increased CBF if tissue PO_2_ can drop. In addition, studies directly measuring tissue PO_2_ indicate that the tissue O_2_ level is reasonably high, about 25 mmHg [49, 50], and until recently the dominant view was that oxygen metabolism is not affected until the tissue PO_2_ is reduced significantly below 1 mmHg [51]. Why does the brain maintain such a high tissue PO_2_ when it seems to be unnecessary, particularly because allowing PO_2_ to drop with increased CMRO_2_ would strongly reduce the need for a large CBF change? More recently, though, the work of Wilson and colleagues [52] has been a strong challenge to previously held views about how oxygen concentration limits metabolism. The basic finding was that the oxygen metabolic rate can be maintained down to very low O_2_ concentrations, but the phosphorylation potential that governs ATP energy metabolism in the cell begins to degrade at a much higher concentration, a partial pressure of about 12 mmHg [53]. This finding is a primary motivation for the thermodynamic framework developed in this paper.

Several other aspects of CBF also are puzzling. Even in the baseline state, OEF in grey matter is relatively low (∼40% [54]), so that the high baseline CBF is already delivering more oxygen than is needed, and OEF drops further with neural stimulation (to ∼30% for a strong neural stimulus [55]). In contrast, heart muscle has a baseline OEF of about 70-80% that does not change much as O_2_ metabolism increases [56], even though the rates of oxidative metabolism are roughly similar for the resting heart and grey matter. Why are the set points and dynamic behavior of blood flow in these two organs so different? Another blood flow effect is that CBF increases strongly with inhaled CO_2_, a phenomenon that has been recognized for more than a century [57], but it is not clear what function this serves.

Finally, the simple picture of serial mechanisms driving the metabolic and flow changes appears to be wrong, or at least a simple path like this is not the primary mechanism involved for CBF control during neural activation. Instead, the current picture is that aspects of neural activity drive CBF changes in a feed-forward way, essentially anticipating the upcoming need for increased metabolism, and this control is applied by a wide variety of mechanisms [58-74]. In this way CBF and CMRO_2_ are driven in parallel by neural activity, and there is growing evidence that the aspects of neural activity that drive the CBF increase may not be the same aspects that entail the largest ATP cost and so account for most of the increased CMRO_2_. As noted above, the primary ATP cost is in restoring ion gradients after neural signaling, primarily excitatory synaptic activity. In contrast, inhibitory synaptic activity often involves opening chloride channels, and because the intracellular/extracellular concentrations are near equilibrium with the resting membrane potential there is little ionic current. As a result, the recovery from inhibitory synaptic activity is likely to be less costly in terms of ATP than the recovery from excitatory activity. Although recovery from spiking of both excitatory and inhibitory neural populations also consumes ATP, estimates for the human brain are that excitatory synaptic activity dominates the overall ATP cost of neural signaling [3]. For this reason, we would expect excitatory activity to be a strong driver of CBF, and a number of studies support this [75]. Interestingly, though, some aspects of inhibitory activity also have a surprisingly strong effect on increasing CBF [62, 65, 66, 76]. The release of nitric oxide (NO), a potent vasodilator, has been associated with the activity of inhibitory neurons [77], and adenosine, which often has an inhibitory neural effect, is also a strong vasodilator [78]. In a recent study using optogenetic methods to stimulate only inhibitory neurons the positive CBF response was approximately the same size as with a more natural stimulus [63]. Why has evolution favored a strong role of inhibitory neural activity to increase CBF?

This complicated picture of how different aspects of neural activity drive CMRO_2_ and CBF suggests the possibility that the balance of changes in CBF and CMRO_2_ may vary with the mix of underlying neural activity [38]. While this would create even more of a problem for any quantitative interpretation of the BOLD signal alone, it potentially opens the door for quantitative physiological fMRI, measuring the changes in CBF and CMRO_2_ [26, 79-88], to provide a deeper and more nuanced interpretation of the underlying neural activity. For this potential to be realized, though, we need a much better understanding of the connections between neural activity, metabolism, and blood flow.

The goal of this paper is to lay a foundation for the theory that the puzzling aspects described above are consistent with thermodynamic limitations on oxygen metabolism [40]. **Section 2** outlines the basic theory, describing the general thermodynamic foundations and how the preservation of the entropy change associated with oxidative metabolism is consistent with the observed physiological behavior. A key result of this development is the importance of maintaining the tissue O_2_/CO_2_ concentration ratio, primarily by modulating CBF. **Section 3** illustrates the implications of preserving tissue O_2_/CO_2_ with modeling studies based on a model of gas transport in blood and tissue, and a simple neural dynamics model of feed-forward drivers of CBF and CMRO_2_, to develop predictions of the theory and compare the predictions with experimental data on CBF responses to neural activity, hypercapnia and hypoxia.

## 2. Proposed thermodynamic framework

### (a) The role of entropy change

The central argument of the thermodynamic framework is that the apparent mismatches of the changes in CMRGlc and CBF compared to CMRO_2_, as well as the CBF response to inhaled CO_2_, can all be viewed as preserving the entropy change of oxidative metabolism in the mitochondria, and through that the oxidative metabolic rate. Entropy is directly related to the number of different molecular states—defined by the positions and velocities of all the particles—which are consistent with given macroscopic constraints. For chemical transformations, such as a chemical reaction or transport of an ion across a cellular membrane, the macroscopic constraint is the average concentrations of different molecules. The basic principles underlying the current theory are that for any transformation of such a system: 1) the net entropy change ΔS of all the processes involved in the transformation cannot be negative (Second Law of Thermodynamics); 2) the net entropy change ΔS has a simple mathematical form that depends on the concentrations of the molecules involved; and 3) the steady-state rate of the chemical transformation depends on both kinetic factors related to the process and on the entropy change ΔS, with that rate going to zero as ΔS goes to zero. All three of these effects follow in a general way from the basic equations of motion of the molecules, specifically that as the molecular states evolve over time they remain as distinct states and do not converge (this is developed in more detail in the *Supplementary Information A: Thermodynamic Basis* based on work in the late twentieth century by E.T. Jaynes [89] and C.H. Bennett [90], and on the implications of the Fluctuation Theorem [91]).

Applying these ideas, for any chemical transformation in a cell the net entropy change cannot be negative. Cellular work involves processes with a negative entropy change, and for these processes to take place they must be coupled to another transformation with a positive entropy change with a larger magnitude. For many cellular processes, the coupled process with a positive entropy change is the breakdown of ATP to ADP and Pi (inorganic phosphate). Another important source of a positive entropy change is the movement of a sodium ion down its electrochemical gradient across the cellular membrane from outside to inside, often used for co-transport of other molecules (e.g., clearance of glutamate from the synaptic cleft by uptake into astrocytes [61]). The sodium gradient is then restored by the sodium/potassium pump, by coupling sodium transport to ATP consumption, and in the brain the activity of the sodium/potassium pump accounts for most of the ATP consumed [11]. The ATP is then restored, involving a negative entropy change, by coupling the process to oxidative metabolism, providing a stronger positive entropy change.

### (b) Key relationships from thermodynamics

The Supplementary Information contains an extended derivation and discussion of the thermodynamic ideas underlying the current theory, leading to the two principal results whose implications are developed below.

#### Entropy change of a chemical process

The first thermodynamic result is the general form of the entropy change ΔS for a chemical transformation. Suppose that a chemical transformation involves the recombination of several reactants (R_1_, R_2_, …) to form several products (P_1_, P_2_,…), and consider one minimal instance of this molecular transformation (e.g., one molecule of R_1_ plus one molecule of R_2_) so that the overall concentrations are not significantly changed. The entropy change associated with this minimal transformation is:

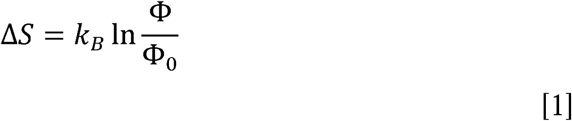

where k_B_ is Boltzmann’s constant and the parameter Φ is the ratio of the reactants to the products of the chemical transformation:

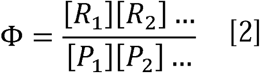

The parameter Φ_0_ is the equilibrium value of Φ such that the chemical transformation involves no change of entropy: ΔS=0. The value of Φ_0_ depends on the specific context in which the chemical transformation occurs, including temperature, the energy change of the chemical system related to the different net binding energies of the reactants and products, environmental interactions such as those between the reactants and products with the surrounding water molecules or with pH, and the conditions under which the chemical transformation takes place (e.g., constant volume or constant pressure). In general, it is difficult to precisely quantify the value of Φ_0_ in a biological setting, but the important result for the current theory is simply the mathematical form of Eq [1].

#### Implications: net entropy change for a linked series of chemical transformations

If a net process contains several steps, such as oxidative metabolism, the net entropy change is the sum of the entropy changes for each step, each of the form of Eq [1]. Mathematically, the net entropy change will then depend on the products of the Φ terms for each step. If the linked steps involve an intermediate chemical that is a product of one step that is consumed as a reactant by the second step, the concentration of that intermediate drops out of the expression for the net entropy change. That is, the concentration of an intermediate will appear in the denominator of Φ_1_ and the numerator of Φ_2_, and the net entropy change will depend on Φ_1_Φ_2_. As a result, for an extended process of chemical transformations, the net entropy change depends only on the concentrations of molecules changed by the net process. However, the concentration of the intermediate can play a useful role in balancing the separate entropy changes of the sequential steps (e.g., increasing the concentration of the intermediate will reduce the entropy change of the first step but increase the entropy change of the second step, without altering the net entropy change), and this idea is further discussed in *Supplementary Information A: Thermodynamic Basis*.

#### Rate of a chemical process

The second important relationship, following from the Fluctuation Theorem (derived in *Supplementary Information A: Thermodynamic Basis*), is that the net rate of a chemical process can be expressed as:

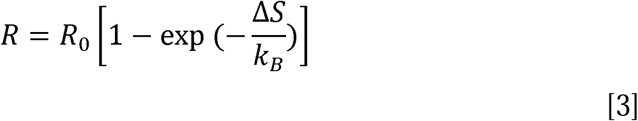

In this form, the rate of a process has a kinetic term *R*_*0*_ and a thermodynamic term depending on the entropy change ΔS. We refer to the parameter *R*_*0*_ as the kinetic rate, and it may depend on the reactant concentrations in a simple way for first order kinetics, or in principle could be independent of the reactant concentrations and controlled by enzyme kinetics (see below). The rate *R*_*0*_ is the rate a process will have if the entropy change is large, and as ΔS goes to zero, the net rate of the process also goes to zero.

#### Implications: Kinetic and thermodynamic effects on the rate of a chemical process

Eq [3] suggests two ways in which the rate of a chemical process could be modified: a kinetic mechanism, changing *R*_0_; or a thermodynamic mechanism, changing ΔS. This distinction is important both for considering how a chemical process can be controlled, and also for considering how a reduction of the concentration of a reactant, such as a reduction of the O_2_ concentration in the mitochondria, can affect the rate of the process through both a kinetic and a thermodynamic limitation. In general, the kinetic term *R*_*0*_ for a process will depend on the enzyme kinetics affecting the mechanics of the process. For example, with simple Michaelis-Menten enzyme kinetics *R*_*0*_ has the form:

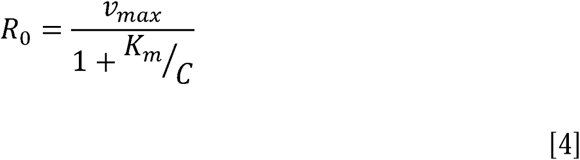

where *C* is the concentration of the reactant, and *v*_*max*_ and *K*_*m*_ are parameters describing the enzyme kinetics. If *C* << *K*_*m*_ the kinetics become first order, and *R*_*0*_ is proportional to *C*. However, for *C* >> *K*_*m*_ the kinetic rate *R*_*0*_ is *v*_max_, independent of *C*. In this case, the rate could be controlled by modulating the enzyme kinetics (i.e., modulating *v*_*max*_), with no change in *C*. However, the net rate of the process is fully determined by those enzyme kinetics only when the entropy change ΔS is large and positive, and as *C* is reduced ΔS also is reduced by Eq [1]. In principle, the reduction of *C* could begin to degrade the rate of the process by a thermodynamic limitation (reduced ΔS) even though the process is not kinetically limited because *C* is still larger than *K*_*m*_.

### (c) Oxidative metabolism in the brain

The oxidative metabolism of Pyr and generation of ATP is a complicated extended process consisting of a series of chemical transformations in which some of the products of one process are the reactants for the next process: 1) in the mitochondria, the TCA cycle metabolizes Pyr, coupled to the conversion of NAD^+^ to NADH and the production of CO_2_; 2) the NADH contributes electrons to the electron transfer chain, which are eventually transferred to O_2_ to produce water, and this electron transfer is coupled to the transport of hydrogen ions across the inner membrane of the mitochondria, creating a proton/potential gradient; and 3) movement of protons down the proton gradient is coupled to conversion of ADP plus Pi to ATP.

For the entropy changes involved, we can consider this extended process as two net processes based on the molecules that are changed (consumed or produced). From Eq [1], the net entropy change associated with the consumption of Pyr and O_2_ and the production of CO_2_ depends on the ratio:

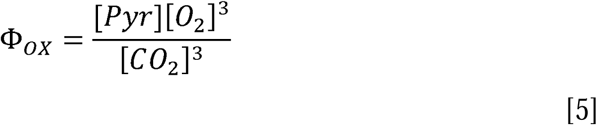

This positive entropy change from oxidative metabolism of pyruvate must be larger than the negative entropy change associated with the conversion ADP+P_i_ ➔ ATP, determined by the phosphorylation potential:

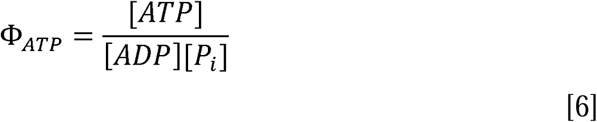

If the tissue O_2_ concentration begins to fall (e.g., by decreased O_2_ delivery), Φ_OX_ will be reduced and the net entropy change ΔS will be reduced. The net positive entropy change can be maintained as Φ_OX_ falls, and CMRO_2_ preserved, if Φ_ATP_ falls as well, as in the study of Wilson and colleagues [53]. In this scenario the metabolic rate is maintained, but at the expense of the entropy change available from ATP to drive cellular work. In the context of the proposed thermodynamic framework, this is interpreted as a distinction between kinetic and thermodynamic limits on metabolism. In this interpretation, the kinetic limit is not reached until [O_2_] drops to a very low level, but the thermodynamic limit is reached at higher O_2_ levels, marked by when Φ_ATP_ begins to degrade to maintain the metabolic rate.

### (d) Preserving the entropy change of oxidative metabolism and the importance of the tissue O_2_/CO_2_ ratio

The core of the proposed thermodynamic framework is that the physiological changes described in **Section 1** serve to preserve Φ_OX_ (Eq [5]) and support the metabolic rate of oxygen. Preserving Φ_OX_ can be done by maintaining the tissue concentrations of O_2_ and Pyr, and the physiological challenge of doing this is the need for transport of these molecules to the mitochondria by diffusion (**Figure 1, lower panel**). Unlike enzymatically controlled processes, where in principle the rate of the process can be controlled by modulation of the enzyme activity with no change in the concentration of metabolic substrates, transport by diffusion necessarily involves concentration gradients. In order to increase the flux of O_2_ to the mitochondria while preserving the mitochondrial O_2_ concentration, the gradient can be increased by raising the O_2_ concentration in blood, and that requires reducing the OEF by increasing CBF more than CMRO_2_. Similarly, to increase the pyruvate flux from the cytosol to the mitochondria, while preserving the mitochondrial pyruvate concentration, the pyruvate gradient can be increased by increasing CMRGlc to increase the cytosolic pyruvate concentration. In this context, the key roles of glycolysis and blood flow are their effects in modulating tissue concentrations of pyruvate and oxygen, respectively, to increase diffusion gradients while maintaining concentrations in the mitochondria. In short, even though empirically it appears that CMRO_2_ may be a lesser player compared with CMRGlc and CBF, because the two latter processes are much more responsive to increased neural activity, within the proposed thermodynamic framework CMRO_2_ is the key function that shapes the complex physiology.

Based on the thermodynamic arguments it is not necessary to preserve the individual concentrations that make up Φ_OX_, but just to preserve Φ_OX_ itself. The role of glycolysis as the first metabolic step, modulating the Pyr concentration, is discussed in detail in the Supplementary Information. In **Section 3(a-f)** we focus on the implications of maintaining Φ_OX_ by maintaining the tissue O_2_/CO_2_ ratio under different conditions, including increased CMRO_2_, hypercapnia, and hypoxia.

## 3. Modeling Results

The key result of the thermodynamic framework is the importance of the tissue O_2_/CO_2_ ratio to preserve Φ_OX_ and thus the entropy change of oxygen metabolism. The modeling studies in this section develop the quantitative implications of this idea, testing the effects of reduced tissue O_2_/CO_2_ and how it can be maintained by blood flow. A quantitative model for O_2_ and CO_2_ transport in blood and exchange with tissue (described in detail in *Supplementary Information B: Oxygen Transport Model*) was used to develop predictions based on the thermodynamic framework and compare the predictions to experimental results. Starting from an assumed normal baseline state, the model predicts the change in tissue O_2_ and CO_2_ concentrations as CBF, CMRO_2_, and arterial blood gases are modulated.

### (a) At what level does reduced tissue O_2_/CO_2_ begin to limit Φ_ATP_?

To put the proposed thermodynamic framework on a more quantitative basis, the limiting O_2_/CO_2_ ratio when the ATP phosphorylation potential Φ_ATP_ begins to degrade was estimated by using the transport model to analyze the extensive data reported by Nioka et al [92] in a study of different degrees of hypoxia in a canine model. Importantly, they measured changes in Φ_ATP_ as well as CBF, CMRO_2_ and blood gases (**Figure 2A**). When analyzed with the transport model, Φ_ATP_ begins to degrade when the tissue O_2_/CO_2_ ratio is reduced to about 50% of the baseline value. **Figure 2B** shows the modeled tissue PO_2_ when Φ_ATP_ begins to degrade, in good agreement with Wilson and colleagues earlier studies in mitochondria preparations finding impairment when tissue PO_2_ drops below about 12 mmHg [52].

**Figure 2.**
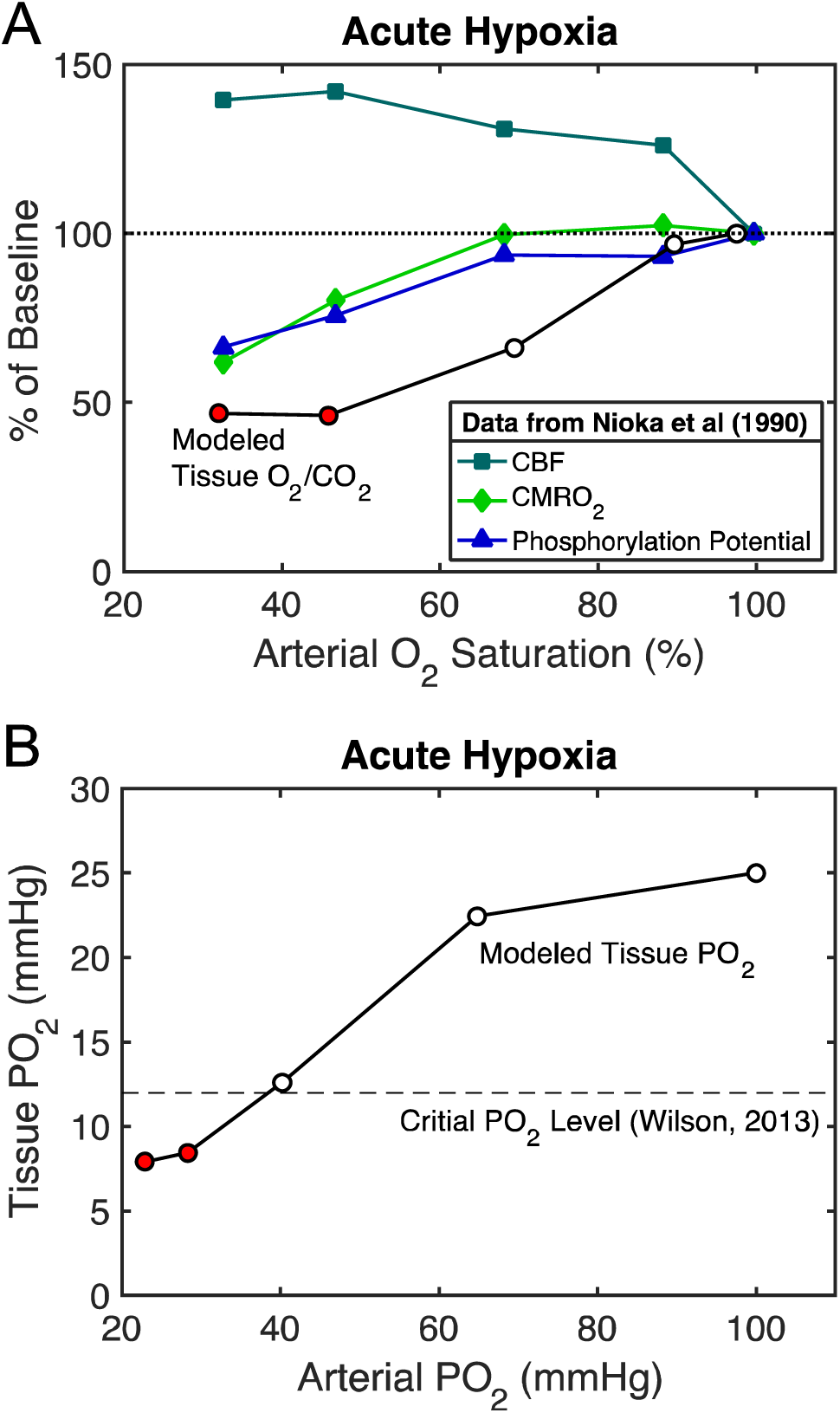
Modeling experimental data to determine the critical level of tissue O_2_/CO_2_. **A**) Data from Nioka et al [92] for several stages of hypoxic hypoxia in a canine model as arterial O_2_-hemoglobin saturation was reduced, including measurement of the ATP phosphorylation potential, are plotted as a percentage of the normoxic value. The transport model was applied to these data to calculate the O_2_/CO_2_ ratio in tissue (circles). The ATP phosphorylation potential was degraded for the two most extreme hypoxic conditions, and for these states the tissue O_2_/CO_2_ ratio was reduced to about 50% of the normoxic baseline value (red circles). **B**) For the same model, the calculated tissue PO_2_ values are plotted as a function of the arterial PO_2_ values. Based on a number of studies of mitochondria, Wilson [52] concluded that a PO_2_ of about 12 mmHg was a critical threshold below which the ATP phosphorylation potential would degrade (dashed line), in good agreement with the data of Nioka et al analyzed with the transport model.

### (b) Effect of reduced O_2_ delivery on tissue O_2_/CO_2_: different sensitivity to reduced flow and reduced arterial PO_2_

Oxygen delivery to the capillary bed depends on both blood flow and the O_2_ content of arterial blood. Because of the nonlinear O_2_-hemoglobin binding curve, with O_2_ delivery largely determined by hemoglobin saturation but O_2_ transport to tissue determined by capillary PO_2_, the degree of impairment of the tissue O_2_/CO_2_ ratio depends on which of the factors affecting O_2_ delivery is reduced. Based on the transport model, the results are that the reduction of tissue O_2_/CO_2_ is more severe when the arterial PO_2_ is reduced than when CBF is reduced (**Figure 3A**). Importantly, when blood PO_2_ is reduced and CBF is increased to restore O_2_ delivery to the baseline level, this is not sufficient to restore the tissue O_2_/CO_2_ ratio (dotted curve in **Figure 3A**). The consequences of reduced arterial PO_2_ and reduced CBF are considered in more detail in the following two sections in the contexts of high-altitude acclimatization and stroke, respectively.

**Figure 3.**
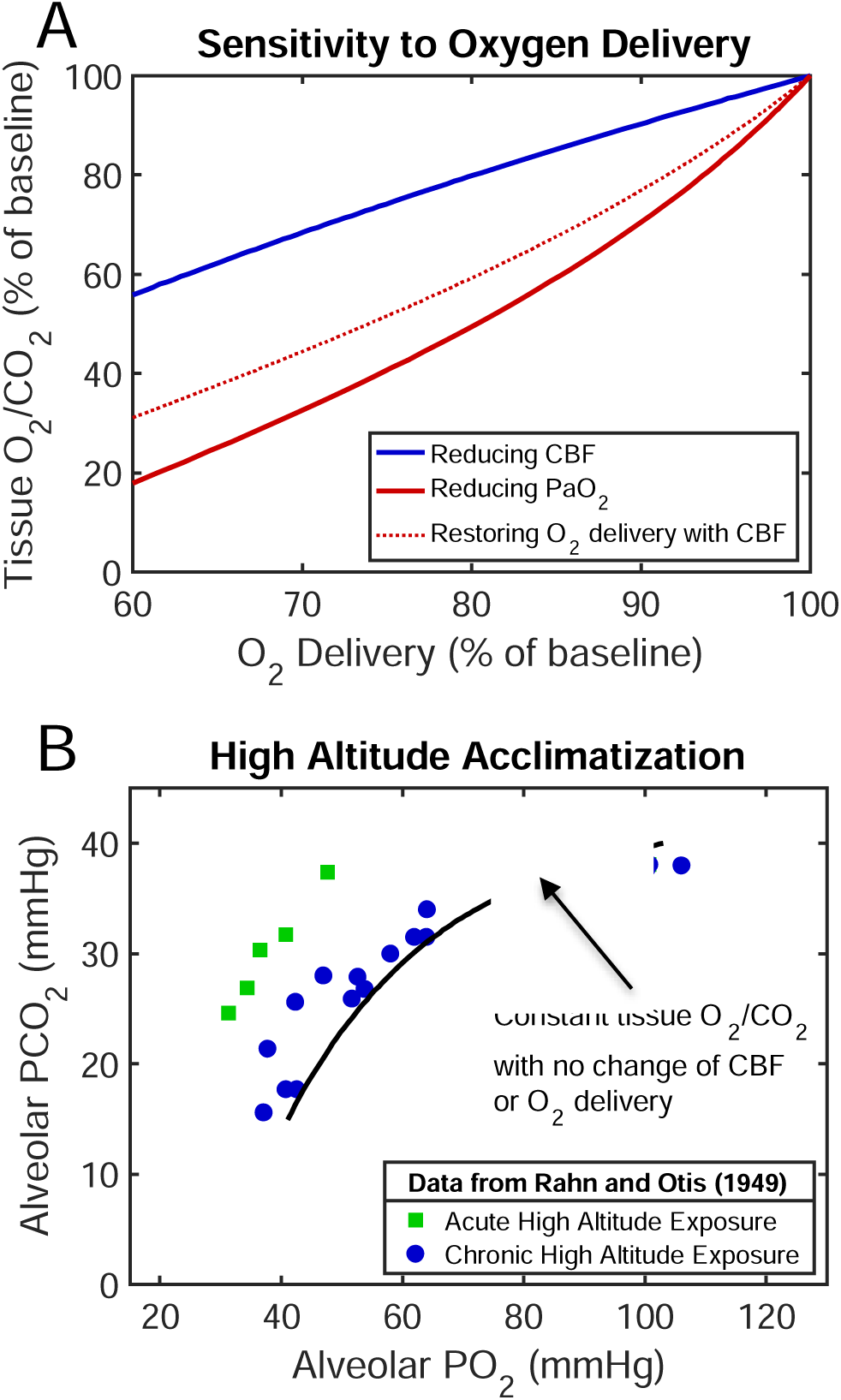
Implications of the thermodynamic framework for sensitivity to oxygen delivery. **A)** Curves for the reduction of tissue O_2_/CO_2_ for two ways of reducing O_2_ delivery: by reducing CBF (blue) or by reducing arterial PO_2_ (red). The brain is more sensitive to reduced O_2_ delivery due to reduced blood PO_2_. Importantly, for the case of reduced arterial PO_2_, restoring baseline O_2_ delivery by increasing CBF does not restore tissue O_2_/CO_2_ (red dotted curve). **B**) Classic data from Rahn and Otis [93] showing the effect of increased ventilation rate as a key element of successful acclimatization to the low inspired O_2_ concentration at high altitude, reducing arterial CO_2_ (PaCO_2_) and increasing arterial O_2_ (PaO_2_), creating a shift diagonally to the right. The solid model curve shows the value of PaCO_2_ needed to compensate for the reduced PaO_2_ and maintain the tissue O_2_/CO_2_ ratio with no change in CMRO_2_ or CBF from the normoxic baseline, and constant O_2_ delivery. (An alveolar/arterial PO_2_ difference of 3 mmHg was assumed in making the plot.)

### (c) High altitude acclimatization: increased ventilation to preserve tissue O_2_/CO_2_ by reducing tissue CO_2_

Long-term exposure to high altitude involves a number of physiological changes, some of which, such as increased hematocrit, serve to restore O_2_ delivery. In addition, though, a key response is an increased ventilation rate that lowers arterial CO_2_, and this response helps to maintain tissue O_2_/CO_2_. For the calculations for the acclimatized state, as arterial PO_2_ was lowered, hematocrit was increased so that O_2_ delivery remained at the baseline level, and there was no change in CBF or CMRO_2_. As in **Figure 3A**, though, maintaining O_2_ delivery, here by increasing the O_2_ carrying capacity of blood, was not sufficient to preserve tissue O_2_/CO_2_. The model was then used to calculate the degree of reduction of arterial PCO_2_ needed in addition to restore tissue O_2_/CO_2_ (**Figure 3B**). Also plotted are the data reported in the classic study of Rahn and Otis [93], reporting alveolar PO_2_ and PCO_2_ for subjects acutely exposed to simulated altitude in a pressure chamber along with values reported from several studies of acclimatized individuals, showing the effect of increased ventilation rate with acclimatization. The model curve is consistent with the experimental data for acclimatized subjects, with the acute subjects falling above the model curve, corresponding to reduced tissue O_2_/CO_2_ if there is no increase of CBF. This too is consistent with other experiments finding increased CBF on acute exposure to high altitude that resolves back to baseline CBF over about a week as subjects acclimatize [94].

### (d) Ischemic stroke: Reducing the fall of tissue O_2_/CO_2_ by reducing CMRO_2_

Calculations with the transport model were used to consider the question of how much of the baseline O_2_ metabolic rate can be maintained as CBF decreases so that the tissue O_2_/CO_2_ level remains above 70% of baseline, which from **Figure 2A** is a level where the phosphorylation potential is still maintained. At this reduced level, and for a normal baseline OEF of 40%, CMRO_2_ can be as high as ∼65% of normal when CBF is reduced to about ∼30% of normal. Interestingly though, for a higher baseline OEF of 60%, the maximum CMRO_2_ is reduced to ∼50% of baseline. Note also that increased OEF by itself is not necessarily a sign of critical impairment: for this example with the normal baseline OEF of 40%, that level of maintained CMRO_2_ is achieved with OEF rising to about 60%. These modeling results suggest that a full evaluation of stroke conditions should involve measurements of both CBF and CMRO_2_, as a more modest reduction of CMRO_2_ than the CBF reduction may be sufficient to maintain tissue O_2_/CO_2_ at a high enough level to preserve the phosphorylation potential.

### (e) Increased CMRO_2_ due to neural activation: CBF response needed to maintain tissue O_2_/CO_2_ while increasing the blood/tissue O_2_ gradient

When CMRO_2_ increases, the gradient of O_2_ concentration between blood and tissue must increase, and this can be accomplished by increasing CBF more than CMRO_2_ so that the O_2_ concentration in blood rises (**Figure 1, lower panel**). Based on the transport model, the CBF required to maintain the tissue O_2_/CO_2_ ratio as CMRO_2_ increases is shown in **Figure 4A**. The CBF/CMRO_2_ coupling ratio *n*, defined as the fractional change in CBF divided by the fractional change in CMRO_2_, varies from about 2 to about 3 as CMRO_2_ increases, in good agreement with experimental studies (reviewed in [40]). Note that the large increase of CBF required is due to the relatively low baseline OEF (40%), and if instead baseline OEF was larger, less of a CBF change would be needed to preserve tissue O_2_/CO_2_.

**Figure 4.**
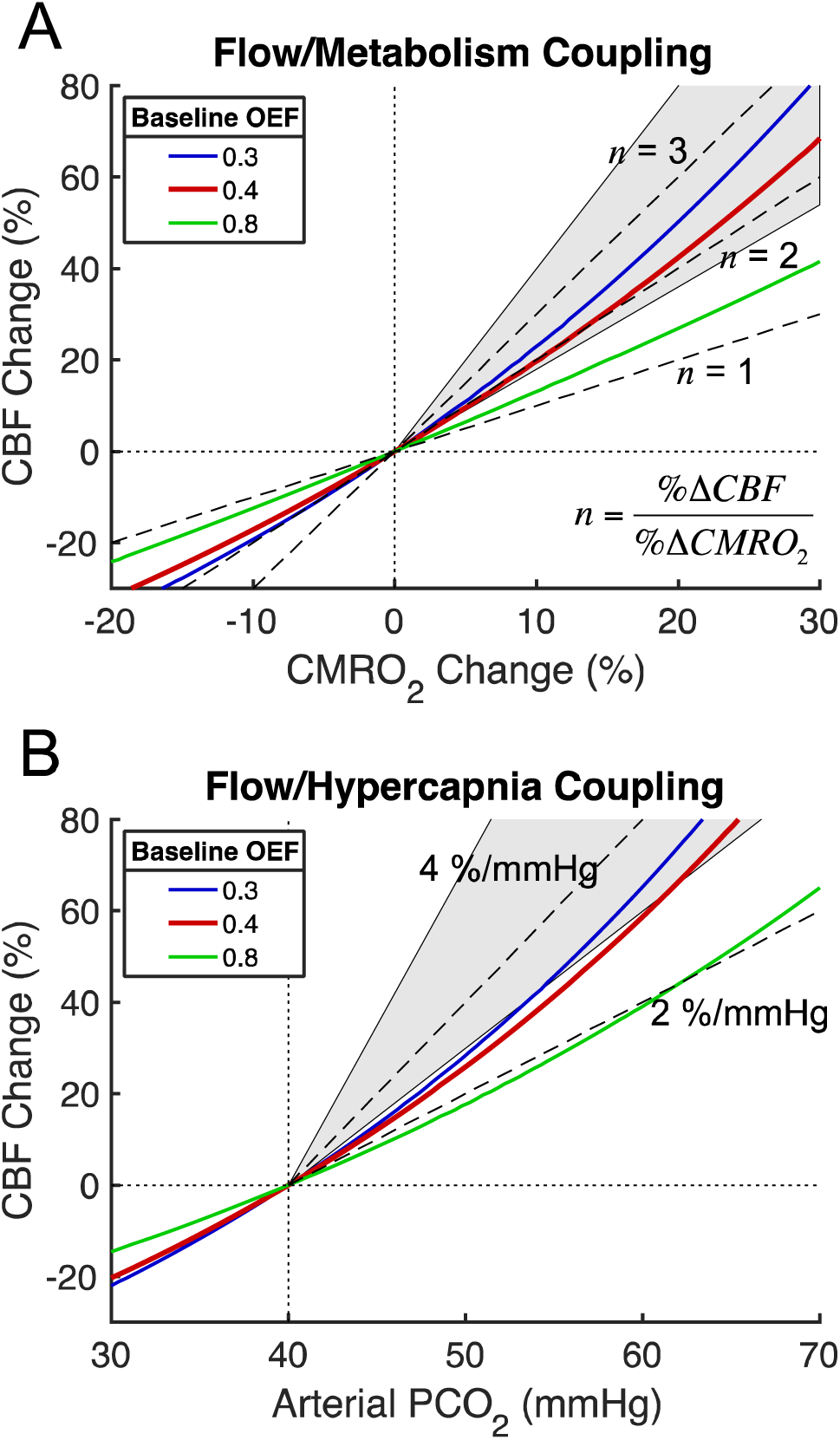
Modeled CBF responses needed to preserve the [O_2_]/[CO_2_] ratio in tissue in response to increased CMRO_2_ and hypercapnia. **A)** With increasing O_2_ metabolism, the required CBF change is shown for three different values of the baseline OEF (the red line is the typical value for brain). Dashed lines show constant coupling ratios *n* of the fractional CBF change to the fractional CMRO_2_ change. The shaded area between *n*=1.8 and *n*=4 shows the approximate range of experimental results. **B)** The required CBF change for increasing arterial CO_2_ shown for the same values of baseline OEF, with dashed lines indicating constant slope values, and the shaded area between slopes of 3 and 7 %/mmHg approximates the range of reported experimental values.

### (f) Increased tissue CO_2_ with hypercapnia: CBF response needed to maintain tissue O_2_/CO_2_ by raising tissue O_2_

Increased CBF, as a mechanism to raise tissue O_2_, also may explain the function served by the strong sensitivity of CBF to hypercapnia. Breathing a gas with increased CO_2_ content will increase arterial and tissue CO_2_ levels, and increased CBF can raise tissue [O_2_] to preserve the tissue O_2_/CO_2_ ratio. **Figure 4B** shows the CBF change needed to maintain tissue O_2_/CO_2_ when the partial pressure of CO_2_ in arterial blood (PaCO_2_) is raised by inhaling a gas mixture with elevated CO_2_. Experimental measurements of the CBF response, expressed as % change per mmHg change in PaCO_2_, vary widely, and the model prediction is consistent with the low end of the experimental results [95-99].

### (g) Neural control of CBF: potential feed-forward mechanisms from both excitatory and inhibitory neural activity

Although the central argument here is that the function served by a large CBF change is to maintain tissue O_2_/CO_2_, there is not yet a known mechanism—an O_2_ sensor—that could be the basis of a feed-back system [40]. Instead, a growing body of evidence points to neural activity, both excitatory and inhibitory, driving the rapid control of CBF in a feed-forward way. In contrast, given the high entropy cost of excitatory activity due to the associated sodium currents and required sodium transport in recovery, we earlier speculated that CMRO_2_ is largely driven by excitatory neuronal activity [38]. This qualitative picture of CMRO_2_ largely determined by excitatory (E) activity with both E and inhibitory (I) activity as feedforward drivers of CBF, was used by Mullinger and colleagues (ref) as a possible explanation for their finding of an altered CBF/CMRO_2_ coupling ratio in a deactivation induced by transcallosal inhibition. In this case the CBF drivers from increased I activity and the associated decreased E activity were in opposition, reducing the net degree of reduction of CBF. A simple example of the reverse effect, when both E and I activity increase—so that the two drivers of CBF change are in the same direction—is the effect of increasing stimulus intensity. Interestingly, an electrophysiology study concluded that while both E and I activity increased as the stimulus intensity increased, there was a growing dispersion between the two with the I activity continuing to increase while excitatory neuronal activity began to taper off [100]. We reported a study in humans [55] measuring BOLD and CBF responses as the contrast of a visual stimulus was increased, finding that the BOLD signal increased more than CBF, which was consistent with a gradual plateauing of the CMRO_2_ response as CBF continued to increase with increasing contrast.

Here we implemented a simple quantitative model as an initial test of the feasibility of these ideas using a version of the Wilson-Cowan (WC) model [101] describing the interaction of excitatory (E) and inhibitory (I) populations of neurons as the stimulus to the E population increases. The model is described in detail in *Supplementary Information C: Neural Model*, but the general structure is illustrated in **Figure 5A**, and the behavior of the E and I populations with increasing input is shown in **Figure 5B.** Assuming that CMRO_2_ is primarily due to E activity and CBF is driven by a combination of E and I activities can create a CBF/CMRO_2_ response curve that is a reasonably good approximation to the curve calculated to be necessary to maintain the tissue O_2_/CO_2_ ratio (**Figure 5C**). The curve is not identical, though, with slight differences in the balance of CBF and CMRO_2_, although these differences have little impact on the general preservation of the entropy change. The data from human visual cortex [55] are also plotted in **Figure 5C**. Interestingly, because the BOLD signal is sensitive to the exact balance of CBF and CMRO_2_ changes, when the BOLD signal is calculated for the WC model curve and for the curve calculated to maintain tissue O_2_/CO_2_ (**Figure 5D**) the WC model gives a much better fit to the experimental data.

**Figure 5.**
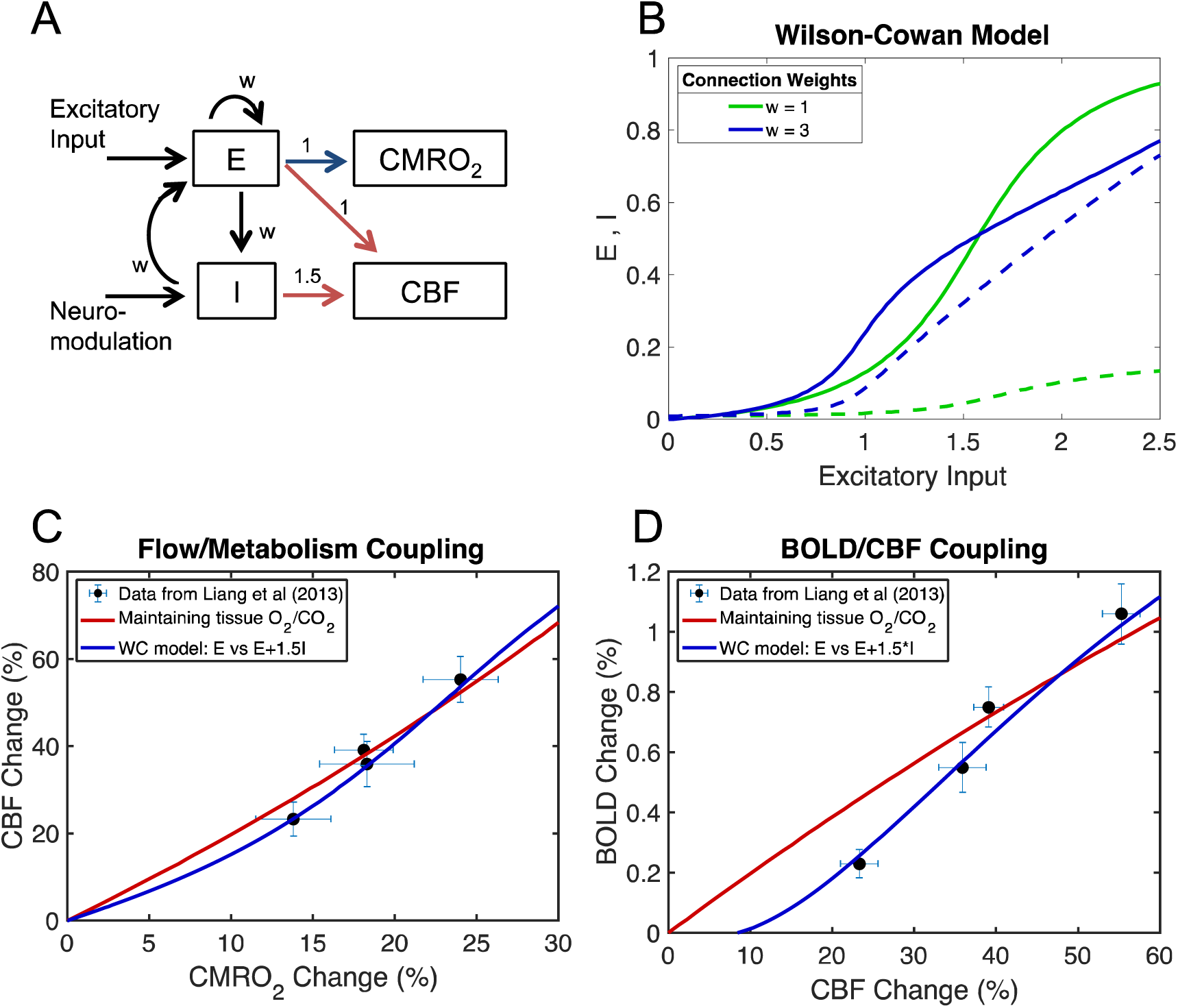
A speculative mechanism for control of CBF through feed-forward signals from neural activity. **A**) The Wilson-Cowan (WC) model [101] was used to model in a simple way the dynamics of interactions of excitatory (E) and inhibitory (I) neural populations driven by an increasing external excitatory input to the E population. The weights for the different interactions were taken to be the same value *w*. As a proof of concept, CMRO_2_ was assumed to be driven by E (blue arrow) and CBF was assumed to be driven by both E and I (red arrows), with a higher weighting for inhibitory activity. **B**) Curves of the activities of the E (solid lines) and I (dashed lines) populations as the input increases are shown for *w*=1 and *w*=3. For higher *w* the E population is more sensitive to weaker input values, with activity rising faster due to the increased self-excitation of the E population, and at higher input values E rises more slowly due to a strong steady rise in I activity. **C**) Data from Liang et al [55] showing the estimated CMRO_2_ changes and the measured CBF changes (mean +/- SEM) from a calibrated BOLD study of effects of increasing contrast of a visual stimulus in humans. The red curve is the curve calculated to preserve tissue O_2_/CO_2_ (from **Figure 2A**). The blue curve is calculated from the WC model with *w*=3 with the assumption that %ΔCMRO_2_ ∼ E and %ΔCBF ∼ E+1.5I, with the same proportionality constant for both chosen to make the largest input value for the curves in panel **B** correspond to %ΔCMRO_2_ = 30%. **D**) Measured BOLD and CBF responses from Liang et al [55] and the predicted BOLD curves for the constant O_2_/CO_2_ model (red) and the WC model (blue). The WC feed-forward model gives a reasonably good approximation of the CBF/CMRO_2_ coupling needed to preserve tissue O_2_/CO_2_ (panel **C**), but the BOLD sensitivity to subtle differences in the balance of CBF and CMRO_2_ creates some divergence of the predictions of the BOLD signal for the two models, with the WC model providing a better fit to the experimental data (panel **D**).

## Discussion

The goal of this work was to lay a foundation for a thermodynamic framework for understanding a number of aspects of blood flow and metabolism in the brain that have emerged from experimental studies over the last several decades. The central implication of the proposed thermodynamic framework is the importance of maintaining the tissue O_2_ to CO_2_ concentration ratio (tissue O_2_/CO_2_) to preserve the entropy change of oxidative metabolism. If this ratio falls, the risk is that the phosphorylation potential also will fall to decrease the entropy cost of converting ADP to ATP. While this could preserve the overall entropy change of oxidative phosphorylation and ATP production, and maintain the oxygen metabolic rate, the critical cost is the reduced entropy change available from ATP that is needed to drive nearly all cellular work. By modeling reported hypoxia data (**Figure 2**), the phosphorylation potential begins to degrade when tissue O_2_/CO_2_ drops to about 50% of its normal value, corresponding to a tissue PO_2_ of about 12 mmHg, in good agreement with the work of Wilson and colleagues (refs).

In the context of maintaining tissue O_2_/CO_2_, blood flow can be thought of as doing more than simply delivering O_2_; it also modulates capillary and tissue PO_2_. The central apparent paradox is that it often appears that blood flow is delivering plenty of O_2_ already, so why does CBF need to increase so much? In the context of the thermodynamic framework, the need for a CBF increase is to meet the central challenge of creating a sufficiently large O_2_ gradient between capillaries and mitochondria to support CMRO_2_ without letting the tissue O_2_/CO_2_ decrease. In response to increased CMRO_2_, the large CBF increase serves to reduce OEF and raise capillary PO_2_, creating the larger O_2_ gradient from blood to tissue needed to support the higher CMRO_2_ while preserving tissue O_2_/CO_2_. In response to hypercapnia, with an associated increase of tissue CO_2_, the CBF increase serves to increase tissue PO_2_ and again preserve tissue O_2_/CO_2_.

In exploring the quantitative implications of maintaining tissue O_2_/CO_2_, the use of a detailed transport model, such as the one used here (*Supplementary Information B: Oxygen Transport Model*), is critical because the results are not intuitively apparent, due primarily to the nonlinear effects of the oxygen/hemoglobin binding curve and the factors that affect it. For example, tissue O_2_/CO_2_ is strongly altered by hypercapnia but much less so by hyperoxia, and it is more sensitive to reduced O_2_ delivery when arterial PO_2_ is reduced than when CBF is reduced. The transport model provides a novel way to evaluate the effect of hypoxia in the brain in different settings by calculating the effect on tissue O_2_/PO_2_, and the limited modeling done here makes predictions related to hypoxia that are in good agreement with experimental data. More work is needed, though, to understand the mechanisms involved and the full implications for high altitude acclimatization, stroke, and other instances of hypoxia.

The modeling also illustrates the way the baseline oxygen extraction fraction (OEF) affects the preservation of tissue O_2_/CO_2_. As noted earlier, a typical OEF in brain is about 40%, and yet for the heart muscle the OEF is typically much higher. At first glance, one might assume that the lower OEF in brain allows for a greater dynamic range of O_2_ metabolism, in some sense a larger buffer of unused capacity that could be accessed by increasing OEF. However, the dynamic range of O_2_ metabolism in the heart is several times larger than in the brain, and with increased O_2_ metabolism in the brain the OEF decreases, rather than increases. The somewhat counterintuitive result of the modeling (**Figure 4A**) is that a given range of blood flow can maintain tissue O_2_/CO_2_ in support of a larger range of O_2_ metabolism (as in the heart) when the baseline OEF is *higher*. However, with a higher baseline OEF the PO_2_ difference between capillary and tissue is reduced if the tissue O_2_ is maintained, and to create the same O_2_ gradient to support O_2_ metabolism the distance from capillary to mitochondrion must be reduced by increasing capillary density. Possibly the relatively low baseline OEF in brain is the result of minimizing capillary density while retaining the capability to maintain tissue O_2_/CO_2_ over a limited range of O_2_ metabolism. In addition, the low baseline OEF in brain confers an advantage in ischemia, and it is here that the “unused capacity” can come into play to preserve a reduced level of CMRO_2_ with increased OEF.

The thermodynamic framework addresses the question of *why* the physiology responds as it does in terms of the function served (i.e., why evolution might preserve these responses), but does not address the question of *how* this is accomplished—the specific mechanisms that drive CBF. Given this picture, with the central role of CBF being to maintain the tissue O_2_/CO_2_ ratio, we might expect that CBF would be strongly driven by an oxygen sensor in tissue, but there is currently no known mechanism that could operate in this way. In addition, O_2_ dissolves so poorly in water that there is very little O_2_ in tissue to serve as a buffer against a drop in O_2_ concentration (we previously estimated that if blood flow stopped completely, the amount of O_2_ dissolved in tissue would support grey matter CMRO_2_ for only about one second, with the O_2_ in trapped blood extending this to about 10 seconds, while in contrast the glucose in tissue provides a strong buffer, able to support normal levels of glycolysis for several minutes [40]). Given the lack of an O_2_ sensor and the lack of a buffer of O_2_, a feed-forward system with a CBF increase triggered by aspects of neural activity could increase O_2_ delivery, and specifically the O_2_ gradient driving O_2_ flux to the mitochondria, in anticipation of the upcoming metabolic need and would help prevent a drop in tissue O_2_ and thus a drop in Φ_OX_. That is, the neural activity initiates sodium fluxes, and afterwards during recovery the sodium will be pumped back at the cost of ATP, and then oxygen metabolism will increase to restore the ATP. As a feed-forward signal, all aspects of neural activity could potentially be beneficial as drivers of CBF, and empirically many mechanisms linking neural activity to CBF have been found [58-74].

As a first test of modeling a neural feed-forward connection as the potential basis of such a mechanism, we assumed that the activities of both inhibitory and excitatory neural populations drive CBF, while CMRO_2_ is driven primarily by ATP consumption related to recovery from the excitatory activity (ref). With this assumption, a simple Wilson-Cowan model of interacting excitatory and inhibitory neural populations was sufficient to explain experimental fMRI data for increasing stimulus amplitude (ref). In this model, the CBF driver from inhibitory activity produced a continuous rise in CBF as stimulus intensity increased even as excitatory activity tapered off, consistent with the data and with the modeled requirement to balance tissue O_2_/CO_2_. We should be cautious about over-interpreting this result, though—it is one experimental data set and one implementation of the model. Nevertheless, it supports the feasibility of modeling the full connections between neural activity and the BOLD signal in a way that is consistent with the proposed thermodynamic framework. Continuing work on possible mechanisms linked to neural activity, both modeling and experimental, and a more detailed understanding of the timing of CBF and CMRO_2_ responses, are needed to determine the full mechanisms involved.

## Supporting information

Thermodynamic Basis

Oxygen Transport Model

Neural Model

## Acknowledgements

The author would like to thank the following for thoughtful comments and suggestions on the ideas of this work: Divya Bolar, Anna Devor, Frank Haist, Susan Hopkins, Frank Powell, G. Kim Prisk, Eulanca Liu, Thomas Liu, Amir Shmuel, Alan Simmons, Aaron Simon, Roger Springett, and Eric Wong.

