## Supplementary material for "The thermodynamics of thinking: connections between neural activity, energy metabolism and blood flow": Thermodynamic Basis

Submitted Nov 1, 2019

Paper for theme issue of *Philosophical Transactions of the Royal Society*, Part B:

'Significant relationships between non-invasive functional neuroimaging and the underlying neuronal activity'

Editors: Clare Howarth, Ralph Freeman, and Anusha Mishra

Corresponding author:
R.B. Buxton
University of California, San Diego, 9500 Gilman Drive, MC 0677, La Jolla, CA 92093-0677, USA

Funding sources:
This work was supported by the National Institutes of Health grants NS036722, MH111359, MH112969, and MH113295.

### Outline of the Supplementary Information in this file

**A. Sketch of thermodynamic ideas underlying the proposed framework**

A1. Thinking about The Second Law

A2. Origin of the Laws of Thermodynamics

A3. Entropy change of a chemical transformation

A4. The Fluctuation Theorem and the rate of chemical processes

A5. Subtleties of linked chemical processes

A6. Glycolysis and oxidative metabolism

A7. Connections with more standard results from equilibrium thermodynamics

**References**

### A. Sketch of thermodynamic ideas underlying the proposed framework

The following is an outline of key thermodynamic ideas to support the proposed thermodynamic framework, specifically the form of the entropy change for a chemical transformation and the effect of the entropy change on the rate of that transformation. The subtlety and complexity of thermodynamics arises because we want to understand the interactions of macroscopic systems, each containing many molecular components, to describe how the world works on a macroscopic scale in the context of appropriate averages over the molecular interactions. The First Law of Thermodynamics, that energy is conserved, is a property of the molecular interactions that carries through to the macroscopic world. The Second Law of Thermodynamics, that entropy does not decrease, is a subtler concept because entropy is not a property of the molecular interactions, but rather a property of the macroscopic constraints that define a macroscopic state and the number of molecular states consistent with those constraints. Importantly, the Second Law is essentially statistical in nature, with an entropy increase more likely than an entropy decrease, and dealing with the associated probabilities in the context of the Fluctuation Theorem is an important part of the development that follows. This development is more of a sketch than a rigorous treatment, designed to motivate the key ideas.

#### A1. Thinking about The Second Law

Entropy and the Second Law of Thermodynamics have always been subtle ideas, and there are ongoing arguments as to their significance and meaning. Here the goal is to thread a path through current ideas about entropy to highlight aspects relevant to understanding brain metabolism, strongly motivated by the work of E.T. Jaynes [1] on the foundations of the Second Law and C.H. Bennett [2] on the thermodynamic limitations on computing. The development here also emphasizes application of the Fluctuation Theorem [3], which was formally developed in the 1990's, to provide a general framework for thinking about the role of entropy change in affecting metabolic rates. That is, although classical thermodynamics focused on equilibrium states, the argument developed here is that the Fluctuation Theorem provides a way to extend these ideas to a nonequilibrium steady-state macroscopic process, such as the metabolic rate of a biological system.

Fundamentally, entropy is a measure of the number of molecular states that are consistent with given macroscopic constraints. In the early development of thermodynamics, motivated by the potential work that could be done by steam engines, macroscopic constraints often took the form of fixed values of parameters such as volume, pressure, and temperature of a gas. For biological applications, the most important macroscopic constraints are the concentrations of molecules involved in transport and chemical reactions, and we are primarily interested in macroscopic average rates of those processes.

The Second Law—that the total entropy change is never negative—is not prescriptive; it does not say what *will* happen in any given circumstance, only that those processes with a net positive entropy change *could* possibly happen. A primary theme in the development of the main text is that for a process to happen the entropy change must be positive and there must be an appropriate kinetic mechanism, and the two together determine the rate of the process. If we imagine a system such as a cell, containing many different kinds of molecules and membranes separating different compartments, a great many chemical reactions and transport processes are possible in principle, even though most of them would have such slow intrinsic reaction rates under biological conditions that they would be negligible over relevant biological time scales. Due to this large range of possible transformations it is difficult to define a meaningful notion of 'entropy' as a property of the system as a whole, or even a clear notion of 'equilibrium', because in principle all of those possible processes would have to be considered. Instead, focusing just on the *entropy change* of a particular transformation (e.g., a chemical reaction or transport process) makes it possible to deal meaningfully with the workings of the cell one process at a time.

The great discovery of evolution is that by making enzymes that catalyze specific reactions, an organism can control which chemical transformations will happen at a significant rate, and of course those catalyzed reactions must conform to the Second Law. As a consequence of the prominence of enzyme controlled processes, the relative entropy changes of two processes in general is not a reliable guide to the relative overall rates of the two processes, because those rates are largely controlled by the enzyme activity. However, a key idea developed in this paper is that the rate of a process is independent of the entropy change (e.g., fully controlled by enzyme activity) only when the entropy change is large, and that as the entropy change reduces to zero the rate of the process also reduces to zero.

In this paper, the important role played by the Second Law in brain physiology derives from this effect of the entropy change on the rate of a biological process. It may be useful, though, to comment briefly on the common linkage of entropy and 'disorder', as this can be misleading. The Second Law is not fundamentally about order and disorder, particularly at the macroscopic scale, although it is often loosely described that way (messy desks and shuffled cards have little to do with physical entropy). Largely due to this mischaracterization, in popular usage the Second Law has taken on more of a metaphorical role in attributing the decay of macroscopic systems to increasing entropy. Entropy increase is often involved, but in a subtle way that is not necessarily related to macroscopic 'order'. For example, consider a bowl that falls off a table and lands on the floor, and consider two scenarios, one in which the bowl breaks into several pieces and one in which the bowl is unbroken: which scenario involves the larger change in entropy? Even though the broken bowl could be considered more 'disordered', the entropy change is larger for the bowl that does *not* break. In both cases the potential energy of the bowl on the table is converted to macroscopic kinetic energy as it falls. When it hits the floor that kinetic energy is dissipated as molecular kinetic energy in the floor and the bowl, and the entropy increase is proportional to the amount of energy dissipated to molecular motions. For the bowl that does not break, all of the original potential energy is dissipated as molecular kinetic energy. However, breaking the bowl requires some of that energy to pull the molecules of the bowl apart (i.e., to overcome the negative binding energy of the forces pulling the molecules together). In short, creating new surface area by breaking the bowl requires energy, so not all of the macroscopic kinetic energy is dissipated as molecular motions, and the entropy change is reduced compared with the intact bowl. Although macroscopic things do tend to fall apart, it is misleading to attribute this to the workings of the Second Law.

In short, to think of entropy change in terms of ‘disorder’, one must adopt a very specific meaning of the word disorder that does not necessarily match with more common usages. Physical entropy is specifically associated with the number of basic molecular states associated with given macroscopic states, and the distinction between different macroscopic states may not involve considerations of entropy. For example, imagine a pool table with the balls neatly organized in a compact motionless triangular arrangement, and the transformation associated with a moving cue ball breaking up that orderly arrangement. The energy of the cue ball is distributed among the other balls in a way that may be hard to predict, depending on the precise angle of impact and energy of the cue ball. The result is that there is an increase of the number of possible macroscopic states, from all of the energy concentrated in the cue ball to that energy being distributed in many possible ways across the other balls. However, this redistribution of energy between macroscopic objects does not involve an entropy change—the number of molecular velocity states of the molecules in a ball doe not change if a constant velocity component in one direction is added to each molecule’s velocity. A physical entropy change only occurs if some of the macroscopic kinetic energy of the balls is dissipated to molecular motions through inelastic collisions or friction.

### A2. Origin of the Laws of Thermodynamics

**Dynamic equations of motion.** The origin of the laws of thermodynamics is in the dynamics of molecular motions and the nature of the interactions between molecules. These interactions can be described by sets of differential equations for the position ($\mathbf{x}_i$) and velocity ($\mathbf{v}_i$) of each particle $i$ with mass $m_i$:

$$\frac{d\mathbf{x}_i}{dt} = \mathbf{v}_i$$

$$\frac{d\mathbf{v}_i}{dt} = -\sum_j \frac{1}{m_i} \nabla U_{ij}(\mathbf{x}_i, \mathbf{x}_j)$$

[A1]

where **x** is a particle's three dimensional position vector **x**=[*x,y,z*] and **v** is the corresponding velocity vector **v**=[$v_x$,$v_y$,$v_z$]. The force acting on particle *i* by particle *j*, that produces an acceleration of particle *i*, is described as the gradient of a potential function $U_{ij}$. The first equation is just the definition of velocity as the time derivative of position. The second equation is essentially Newton's second law for a force (*f*=*ma*, *a*=acceleration). The remarkable feature of these equations is that the acceleration of a particle depends only on the positions of all the particles, and not on their velocities. This restricted form of the basic equations of motion leads to the laws of thermodynamics that govern the behavior of macroscopic systems.

**Energy change.** The first consequence of this mathematical form of the equations is that a particular function of the positions and velocities, defined as the total energy *E*, remains constant as the system evolves over time:

$$E = \sum_i \frac{1}{2} m_i \mathbf{v} \cdot \mathbf{v} + \sum_{i \neq j} \frac{1}{2} U_{ij}$$

[A2]

with the kinetic energy represented by the first term and the energy associated with the potential that defines the force, the potential energy, represented by the second term. When macroscopic systems interact, the energy of each system may change but the sum of the energy changes is always zero. Because of this constancy, we often speak of an "energy flow" from one system to another, as the energy of one system goes down and the energy of a second system goes up by the same amount, as though some kind of substance moved between them and the amount of that substance is fixed. It is hard to resist thinking of it this way, as analogous to the way we might think of conservation of the number of particles that does not change as the particles are rearranged. But it is good to remember that formally the conservation of energy is a mathematical property, rather than the idea that energy exists as some kind of “substance”. For example, energy can take on negative values, so the conservation of energy has a different character than the conservation of matter.

**Entropy change.** A macroscopic system, containing many particles, can be characterized by the macroscopic constraints that are imposed on the molecular states, such as a fixed total volume, total number of particles, and total energy. The entropy $S$ associated with a macrostate is defined by Boltzmann’s equation in terms of the total number of molecular states $W$ that are consistent with the associated macroscopic constraints: $S = k_B \ln W$, and the constant $k_B$ is Boltzmann’s constant. Each one of those states $W$ can be fully described by a list of all the $3N$ position components and $3N$ velocity components of the $N$ particles at a particular time for that state. By considering a $6N$ dimensional phase space, in which each axis corresponds to a single position or velocity component of a particle, the set of numbers describing a molecular state can be represented as a single point in phase space. A given macroscopic state is then associated with a set of points in phase space, and the entropy of that macroscopic state is a measure of the number of those points. Absolute entropy, though, never enters into the physics of macroscopic processes. Instead, the key factor involved in the physics is the change in entropy $\Delta S$ associated with the transformation from a macroscopic state $A$, consistent with $W_A$ molecular states, to a macroscopic state $B$, consistent with $W_B$ molecular states:

$$\Delta S_{A \to B} = k_B \ln \frac{W_B}{W_A} \quad \text{[A3]}$$

**Motion reversal symmetry.** Like the First Law of Thermodynamics (energy conservation), the Second Law also has its origin in the mathematical form of the basic equations of motion. Each of the $W$ different molecular states consistent with a given macrostate is defined by distinct combinations of particle positions and velocities at a time $t$=0. Over time, each state evolves according to the dynamical equations of motion, with changing particle positions and velocities, with each state tracing out a trajectory in phase space. The crucial consequence of the equations of motion is that the states remain distinct at all future times. That is, at no time do two of the original states ever converge on the same point in phase space. The reason is due to a reversible feature of the equations of motion. After a state has evolved for a time $t$, if we imagine reversing each of the velocities and then allowing the system to continue to evolve, the system will return to the original configuration of positions with each of the velocities reversed. If two distinct initial states ever evolved to a time where they occupied the same point in phase space, the motion reversal symmetry would fail since that single point could not evolve back into two distinct states after motion reversal. (More formally, in statistical mechanics the equations of motion are expressed by Hamilton's equations, and by Liouville's theorem the probability density in phase space is conserved.) The motion reversal symmetry holds because the equation for the force on a particle depends only on the positions of the particles, and not on their velocities. If a term proportional to $-\boldsymbol{v}$ was added to the right side of Eq [A1], the motion reversal symmetry would no longer hold (nor would energy conservation). At the macroscopic scale, 'dissapative' systems do occur, in which the macroscopic energy decays away, and these systems are modeled in a phenomenological way with a resistance or relaxation term. However, the full picture is that the macroscopic energy disappears but the kinetic energy of the molecular motions increases so that the net change in energy is zero.

**The Second Law**. Now consider the transformation from a macroscopic state $A$ to a different macroscopic state $B$, defined by a different set of macroscopic constraints. If each of the molecular states $W_A$ associated with macrostate $A$ reliably evolves into molecular states consistent with macrostate $B$, then the entropy of state $B$ cannot be less than the entropy of state $A$ because all those original states are still distinct from one another. It may be, though, that the macroscopic constraints of state $B$ are consistent with a larger number of molecular states $W_B$ than the original state $A$, and in that case the entropy would increase in the transformation from state $A$ to $B$. In short, for any transformation from a macrostate *A* to a macrostate *B* that happens reliably (i.e., all of the molecular states that are initially consistent with *A* evolve to a configuration consistent with *B*) the entropy must increase or stay the same. This is essentially Jaynes explanation of the origin of the Second Law as a source of well-defined consistent behavior of macroscopic systems [1].

**A3. Entropy change of a chemical transformation**

**Counting states**. The mathematical form of the entropy change is important, because it allows the entropy change to be a well-defined physical quantity even when the entropy itself is hard to define. For example, consider a gas of $N$ particles in a volume $V$: what is the entropy change associated with a change of the macroscopic constraint from a volume $V_1$ to a volume $V_2$? If we start with a volume $V$ and ask how many different positions a particle could have, the answer is infinite if position is a continuous variable, so the entropy itself is not well-defined. However, we can approach a calculation of the entropy change by imagining the volume $V$ to be composed of a finite number of 'minimal volumes' $\delta V$, such that each particle could be in one of the finite $V/\delta V$ volume elements (positions). The number of ways of distributing $N$ particles across these volume elements is then $(V/\delta V)^N$. The entropy for a macroscopic state with volume $V$ is still ill-defined because of the artificial introduction of $\delta V$, but if we focus just on the entropy change the $\delta V$ divides out, leaving a well-defined expression for the entropy change: $\Delta S = k_B \ln(V_2/V_1)$. Note that the introduction of $\delta V$ is artificial from a classical physics view, but from a quantum physics view there is a minimum granularity of phase space due to the uncertainty principle, measured in terms of $h^3$, where $h$ is Planck's constant.

**Entropy change for diffusion across a membrane**. As a simple example, consider two compartments (designated 'left' and 'right', with respective volumes $V_L$ and $V_R$) separated by a membrane with channels that allow a molecule to pass between the compartments, and define a macroscopic state *A* as having $N_1$ molecules in the left compartment and $N_2$ in the right compartment, and a macrostate *B* as having $N_1-1$ molecules in the left compartment and $N_2+1$ molecules in the second compartment. The transformation from state *A* to state *B* is then the movement of one molecule from left to right across the membrane and our goal is to determine the associated entropy change. The number of states $W_A$ consistent with macrostate *A* depends on the product of two factors: 1) the number of ways *N* particles ($N=N_1+N_2$) can be distributed between the two compartments such that there are $N_1$ particles on the left and $N_2$ particles on the right, which is $N!/(N_1!N_2!)$; and 2) the number of position states possible for $N_1$ particles in volume $V_L$ and $N_2$ particles in volume $V_R$, which, following the approach above, can be taken as $(V_L/\delta V)^{N1}(V_R/\delta V)^{N2}$. A similar expression applies to $W_B$, but with $N_1$ decreased by one and $N_2$ increased by one. The entropy change in going from macrostate *A* to macrostate *B* is then:

$$\Delta S = k_B \ln \frac{N_1}{N_2 + 1} \frac{V_R}{V_L} \approx k_B \ln \frac{C_1}{C_2} \qquad [A4]$$

where $C_1$ and $C_2$ are the concentrations of particles in the left and right compartments, respectively. The entropy change is positive when $C_1>C_2$, negative when $C_2>C_1$, and equal to zero when $C_1=C_2$.

**Entropy change of a chemical reaction**. The entropy change for the diffusion case is an exceptionally simple example, and for most transformations it is difficult to define the full set of changes that alter the number of states. However, we can develop a general expression for the entropy change of a chemical reaction without knowing the full details of the transformation. Consider a chemical reaction in which a molecule XY separates into a molecule X and a molecule Y. This also will involve changes in the surrounding bath of molecules, including an energy change needed to balance the energy change of the chemical system related to the binding energy of the XY molecule, interactions between the molecule and the bath of surrounding molecules (mostly water), and possibly a volume change depending on the conditions under which the chemical transformation takes place (e.g., constant volume or constant pressure). Importantly, though, the number of states consistent with a given macroscopic constraint of concentrations [XY], [X] and

[Y] can be taken as $W=W_0W_C$, where $W_0$ includes all of the unknown factors related to interactions with the bath of molecules, and $W_C$ simply describes the number of ways of creating the concentrations defining the macroscopic state with a fixed number of total X and Y molecules (i.e., including the bound ones in XY). The transformation of one molecule of XY→X+Y then involves a change $W_0 \rightarrow W_0'$ and $W_C \rightarrow W_C'$, so the entropy change is:

$$\Delta S = k_B \ln \frac{W_0'}{W_0}\frac{W_C'}{W_C} = k_B \ln \frac{W_0'}{W_0} + k_B \ln \frac{W_C'}{W_C} = \Delta S_0 + \Delta S_C$$

[A5]

The entropy change $\Delta S_C$ due just to the number of configurations of the molecules is additive to all the other sources of entropy change $\Delta S_0$. The number of states $W_C$ corresponding to the chemical configuration is the number of ways $N_{X0}$ total X molecules and $N_{Y0}$ total Y molecules can be arranged to form $N_{XY}$ bound molecules and $N_X$ and $N_Y$ free molecules is:

$$W = \frac{N_{X0}!}{(N_{X0} - N_{XY})!}\frac{N_{Y0}!}{(N_{Y0} - N_{XY})!}\frac{1}{N_{XY}!}$$

[A6]

For the transformation of one XY molecule, $N_{XY}$ is reduced by one and $N_X$ and $N_Y$ are each increased by one, so the entropy change due to the configuration change is:

$$\Delta S_C = k_B \ln \left( N_{XY} \frac{1}{N_{X0} - N_{XY} + 1}\frac{1}{N_{Y0} - N_{XY} + 1} \right) \approx k_B \ln \frac{N_{XY}}{N_X N_Y}$$

[A7]

We now assume that for the full set of factors that define $\Delta S_0$, there is an equilibrium state in which the transformation of one molecule XY→X+Y is in equilibrium, such that $\Delta S_0 = -\Delta S_C$. That is, there is a particular configuration of the concentration ratios that would be in equilibrium with all of the factors (temperature, bond energy changes, etc) such that the system is in equilibrium with regard to this particular transformation. Then from Eq [A5} the entropy change for any different configuration of the molecules can be expressed as:

$$\Delta S = k_B \ln \frac{\Phi}{\Phi_0}$$

[A8]

where $\Phi$ is the ratio [XY]/[X][Y], and $\Phi_0$ is the value of that ratio when the transformation XY→X+Y is at equilibrium. This form of ΔS can be generalized to any process in which a set of reactant molecules $R_1$, $R_2$, etc are recombined into a set of product molecules $P_1$, $P_2$, etc with:

$$\Phi = \frac{[R_1][R_2] \ldots}{[P_1][P_2] \ldots}$$

[A9]

**A4. The Fluctuation Theorem and the rate of chemical processes**

Returning to the motion reversal symmetry of the basic dynamic equations and the origin of the Second Law, we can take this argument one step farther and bring in the idea of the Fluctuation Theorem. The argument is general, but for concreteness consider the diffusion example described above. The number of molecular states consistent with our two macroscopic states *A* and *B* are $W_A$ and $W_B$, respectively, at time $t$=0. We assume that for each state consistent with *A*, if we reverse all the velocities that new state is also consistent with *A* (i.e., that new state is another of the original $W_A$ states), and similarly for state *B*. If the states consistent with *A* now evolve for a time interval τ, some of those states will evolve such that a molecule on the left encounters a channel and passes through to the right side, a transition from macrostate *A* to macrostate *B*. If the number of states that make this transition is $W_{A \rightarrow B}$, the probability of macrostate *A* transforming to state *B* during the time τ is then the probability that the actual molecular state is one of these states, $p_+ = W_{A\rightarrow B} / W_A$. Now consider the reverse transformation, from *B* to *A*, involving the movement of one molecule from the right compartment to the left: how many molecular states $W_{B\rightarrow A}$, initially consistent with macrostate *B*, will make the transition to state *A* during the time interval τ? We can construct a set of molecular states that will transition from *B* to *A* by letting each of the $W_{A\rightarrow B}$ molecular states initially consistent with *A* evolve for a time τ and then reversing the velocities of each state. Each of these new states is initially consistent with macrostate *B* and will evolve to be consistent with macrostate *A* during an interval τ. In fact, there can be no additional molecular states consistent with *B* that would transition to *A*, because again by the motion reversal symmetry, such a state would have to correspond to one of the original $W_{A\rightarrow B}$ states. The key consequence of the motion reversal symmetry is thus that $W_{B\rightarrow A} = W_{A\rightarrow B}$, and the probability of the reverse transition happening during time τ is $p\_ = W_{B\rightarrow A} / W_B$. This leads to the basic statement of the Fluctuation Theorem: for two macrostates *A* and *B*, the probability of the forward transition from *A*→*B* during a time interval τ and the probability of the reverse transition from *B*→*A* are directly related to the entropy change $\Delta S_{A\rightarrow B}$ between the two states:

$$\frac{p_+(\tau)}{p_-(\tau)} = \exp\left(\Delta S_{A\rightarrow B}/k_B\right)$$

[A10]

**The rate of a process**. Building on the Fluctuation Theorem, the net rate of a process is:

$$R = \frac{p_+(\tau)}{\tau}\left[1 - \exp\left(-\frac{\Delta S}{k_B}\right)\right] = R_0\left[1 - \exp\left(-\frac{\Delta S}{k_B}\right)\right]$$

[A11]

where ΔS is the entropy change of the forward direction. In this form, the rate of a process has a kinetic term $R_0$ and a thermodynamic term depending on the entropy change ΔS. In the argument above, the details of exactly what determines $R_0$ (i.e., the mechanics that determines $W_{A \to B}$) did not have to be specified to demonstrate the role played by the entropy change. The rate $R_0$ is the rate a process will have if the entropy change is large, and as ΔS goes to zero, the net rate of the process also goes to zero.

**Diffusion example revisited.** As a simple example of the implications of Eq [A11], we return to the earlier example of diffusion between two compartments. For the simplest case of diffusion of an uncharged molecule across a membrane the equilibrium ratio is $\Phi_0=1$. (For an ion moving across a membrane with an electric potential across it, the equilibrium ratio will differ from one depending on the voltage across the membrane.) From Eq [A4], the entropy change is $\Delta S = k_B \ln(C_1/C_2)$. The kinetic term in Eq [A11] is of the form $R_0 = \kappa C_1$, the unidirectional flux from the left compartment to the right compartment, where κ is proportional to the product of the membrane permeability and surface area. Combining these expressions for the kinetic term $R_0$ and the thermodynamic term ΔS in the general rate equation [A11] gives: $R = \kappa(C_1\text{-}C_2)$, the familiar Fick equation for diffusion across a membrane. When the entropy change goes to zero ($C_1 = C_2$), the net rate of the transformation (the net rate of transfer of particles from the left to right compartments) also goes to zero.

**Producing RNA matched to DNA**. Bennett [2] used Eq [A11] to describe the activity of RNA polymerase constructing an RNA molecule from a DNA template. The argument was that the process of attaching a base to the growing RNA molecule is reversible, and in principle construction of the RNA could occur with no entropy cost if the system could tolerate the extremely slow rate of construction. By coupling the addition of a base to the conversion of an ATP molecule, the entropy change increases and so does the rate of the process. This argument was developed in the context of investigating the thermodynamic limits to computation, with this as a biological example with similarities to reversible computing.

**Linked processes.** For a sustained linked process, such as the steady-state metabolism of Pyr and $O_2$ to generate ATP, the rates for each stage are the same and each depends on the associated kinetic term and entropy change for that stage. For example, consider a simple two-step process in which the product of the first step is the reactant for the second step. For a steady-state rate R, the requirement is:

$$R = R_1 \left[1 - \exp\left(-\frac{\Delta S_1}{k_B}\right)\right] = R_2 \left[1 - \exp\left(-\frac{\Delta S_2}{k_B}\right)\right]$$

[A12]

The rate $R$ could be controlled either by changing the individual kinetic terms ($R_1$ and $R_2$) or by changing the individual entropy changes at each step. Again, in principle the net rate $R$ could be controlled just by the kinetic rates $R_1$ and $R_2$ associated with the two steps, as long as the entropy change of each process is maintained.

**A5. Subtleties of linked chemical processes**

Oxidative metabolism is a chain of chemical processes involving chemical reactions and transport of molecules across membranes. As a simple example of linked processes, consider a two-step process in which X→I followed by I→Y, where X is the initial molecule, Y is the final molecule and I is an intermediate molecule. Each step has an associated entropy change ΔS given by Eq [1], with $\Delta S_1$ of the first step depending on [X]/[I] and $\Delta S_2$ of the second step depending on [I]/[Y]. The net entropy change of the extended process is the sum of these individual entropy changes, and due to the mathematical form of Eq [1], the net entropy change depends just on [X]/[Y] with the dependence on [I] dropping out. In short, for a chain of reactions the net entropy change depends only on the concentrations that have a net change due to the process, with any intermediate compounds that are created and then consumed during the extended process having no contribution.

In general, though, the Second Law requires that the entropy change ΔS at each step of an extended process must be positive, not just that the net entropy change of the entire extended process is positive. For a simple two step process, in which the intermediate product I of the first step is the reactant for the second step, we need to distinguish two scenarios. If the intermediate molecule I is fed directly to the second step, the two steps are tightly coupled and the net entropy change is what must be positive. For example, the coupling of ATP hydrolysis to ion transport by the sodium/potassium pump is an example of tightly linked processes, and the net entropy change is the sum of the entropy changes for the change in ATP and for the ion transport. However, if the first step feeds I molecules to an intermediate pool, and the second step then draws from this intermediate pool, the requirement of the Second Law is that both $\Delta S_1$ and $\Delta S_2$ must be positive. For example, metabolism of pyruvate in the mitochondria adds to the pool of NADH, and NADH is then a reactant for the oxygen metabolism step.

For our simple X/Y example, a reduction of [X] would reduce both the net entropy change and the entropy change of the first step, $\Delta S_1$. A parallel reduction in [Y] would preserve the net entropy change, by increasing $\Delta S_2$, but by itself would not compensate for the reduction of $\Delta S_1$. If $\Delta S_1$ is reduced to zero the net process cannot go forward even though the net entropy change is positive, because X and Y are involved in different reactions. However, the possibility of adjusting the concentrations of intermediates provides a way to balance the entropy changes across steps of the net process while keeping the net $\Delta S$ constant. In this example, a parallel reduction in [I] (in addition to [Y]) would preserve $\Delta S_1$ as well as $\Delta S$.

In addition, the adjustment of the concentration of intermediates can happen relatively automatically. Imagine that the simple X→Y system above is embedded in a larger chain of processes involving the creation or delivery of X and the later consumption or clearance of Y. If this process is in a steady-state, then the two steps have the same net rate, each of the form of Eq [A12], with $R_1$ and $\Delta S_1$ for the first step and $R_2$ and $\Delta S_2$ for the second step. If now the concentration of X is reduced, the production of the intermediate I will drop, and as step 2 proceeds at the original steady-state rate, the concentration of I will drop. This, in turn, will begin to decrease the rate of the second step, and thus decrease the production of Y, and the concentration of Y will be reduced. In principle, this process of transient adjustment can lead to recovery of the original steady-state rate with the original entropy changes of the two steps, but with lower individual concentrations of X, Y and I.

In short, the arguments above illustrate two important principles for a steady-state of linked chemical processes in which the products of one step add to a metabolite pool that serves as the substrate for the next step: 1) the net entropy change depends only on the concentrations of the inputs and final outputs, and not on the concentration of intermediate molecules that are created and then consumed; and 2) transient adjustments of the intermediate concentrations can serve to maintain the overall flux and entropy change as the concentration of reactants changes. Both effects are important in oxidative metabolism.

**A6. Glycolysis and oxidative metabolism**

As discussed in the main text, by preserving the overall entropy change, and adjusting intermediate concentrations to balance the entropy changes of the stages, the net metabolic rate can be maintained for a linked set of chemical transformations. In a simplified form we can think of the metabolism of glucose and $O_2$ as happening in two connected stages: 1) the breakdown of glucose into pyruvate in the cytosol by glycolysis and then the further breakdown of pyruvate to $CO_2$ in the TCA cycle in the mitochondria with the transfer of electrons from $NAD^+$ to NADH; and 2) the donation of electrons from NADH to the electron transfer chain (ETC) with final acceptance of the electrons by $O_2$ to form water. For the first stage, the entropy change $\Delta S_1$ associated with the metabolism of pyruvate in the TCA cycle increases with increased [Pyr]/[$CO_2$] and [$NAD^+$]/[NADH] ratios, and the entropy change $\Delta S_2$ of the ETC increases with [$O_2$] and the [NADH]/$NAD^+$] ratio. Glycolysis in the cytosol is a physiological mechanism well-positioned to both maintain or increase the entropy change $\Delta S_1$ of the first stage by modulating $[Pyr]_{mit}$, the pyruvate concentration in the mitochondria, and to additionally increase the concentration of the intermediate NADH to help increase the entropy change $\Delta S_2$ of the second stage, as described below.

Pyruvate and NADH are both created by glycolysis in the cytosol at a rate determined by CMRGlc, and there are then two different fates for these molecules. First, the pyruvate can be converted to lactate with the conversion of NADH back to $NAD^+$, and the lactate cleared from the cell. For the cell, this fate contributes to ATP generation only through the production of two ATP in glycolysis (although if the lactate shuttle idea is operating the lactate could be taken up by another cell and further metabolized). The second fate is that the pyruvate diffuses into the mitochondria where it is metabolized at a rate proportional to $CMRO_2$. The rate of entry of Pyr into the mitochondria can be modeled as proportional to a concentration gradient $[Pyr]_{cyt} - [Pyr]_{mit}$ between average concentrations in the cytosol and the mitochondria. If $CMRO_2$ increases, the rate of Pyr delivery to the mitochondria must increase in proportion by increasing that gradient. To increase the Pyr gradient while preserving $[Pyr]_{mit}$, the $[Pyr]_{cyt}$ must be increased. This requires CMRGlc to increase more than $CMRO_2$. In addition, there is also a mechanism for increased [NADH] in the cytosol to be converted to increased [NADH] in the mitochondria. That is, the NADH molecule cannot pass through the mitochondrial membrane, but the malate/aspartate shuttle is a mechanism for transferring the electrons carried by NADH to the shuttle, with the change of cytosolic NADH to $NAD^+$, and then the transfer of the electrons from the shuttle to $NAD^+$ in the mitochondria, creating NADH. In short, the mechanism of increasing CMRGlc more than $CMRO_2$ can serve to maintain or increase mitochondrial [Pyr], and thus maintain $\Delta S_1$, and also increase mitochondrial NADH, benefitting $\Delta S_2$.

**A7. Connections with more standard results from equilibrium thermodynamics**

For the development of the thermodynamic framework in the main article the two key results from thermodynamics are the mathematical form of the entropy change for a chemical process (Eq [A8]) and how the entropy change affects the rate of the process (Eq [A11]). The derivation sketched above was based on physical principles of entropy to emphasize the generality of the results. However, this approach essentially bypassed many of the standard elements familiar in thermodynamics, such as temperature, work and free energy. In this section these concepts are considered in light of the previous arguments, and the connections with the form of the entropy change (Eq [A8]) are discussed.

**Temperature.** In classical thermodynamics, if energy ΔE is added to a system the change in entropy ΔS is:

$$\Delta S = \frac{\Delta E}{T}$$

[A13]

where T is the temperature of the system (from the work of Kelvin and Clausius). In fact, one could take this as the definition of temperature, as the proportionality between energy change and entropy change. In this role T has the familiar features of temperature. If energy ΔE is transferred from a system with temperature $T_1$ to a system with temperature $T_2$, the net entropy change is $\Delta S = -\Delta E/T_1 + \Delta E/T_2$. That entropy change is positive only if $T_1 > T_2$, so a spontaneous energy transfer can only happen from a hotter to a colder system. If $T_1 = T_2$ the two systems are in equilibrium with respect to energy transfer ($\Delta S = 0$).

Eq [A13] also follows from Boltzmann's equation [A3]. Consider an ideal gas of N particles that do not interact with each other, so that the energy of the system is all in the kinetic energy of the particles. As energy is added to the system the number of available momentum states will increase, increasing the entropy. Assuming the volume does not change, so there is no change in position states, we can just consider the 3N-dimensional momentum space component of phase space, with each axis corresponding to one momentum component of one particle. The total energy E of the system defines the radius $r$ of a hypersphere in this space, with $r$ given by $r^2 = 2mE$ (i.e., the kinetic energy associated with a particle with momentum p is $p^2/2m$). The number of momentum states W consistent with total energy E is proportional to the surface area of this hypersphere: $W \sim r^{(3N-1)}$. When E increases by a small amount ΔE, the entropy change is then:

$$\Delta S = k_B \ln \frac{W(E+\Delta E)}{W(E)} \approx k_B \ln\left(1+\frac{\Delta E}{E}\right)^{3N/2} \approx k_B \frac{3N}{2E}\Delta E$$

[A14]

where the approximations are valid for N>>1 and ΔE/E<<1. For consistency with classical thermodynamics and Eq [A13], the factor multiplying ΔE is then 1/T. In addition to defining temperature as the proportionality between ΔE and ΔS, this derivation shows a new feature of temperature: the average energy associated with a single momentum component of one particle (E/3N) is $k_B T/2$. The factor $k_B T$ thus reflects the scale of the individual particle energies.

In short, the temperature T plays two roles: $k_B T$ is the scale of kinetic energy associated with one component of the momentum of a particle, and 1/T is the proportionality between an amount ΔE of energy dissipated in the thermal bath and the resulting change of entropy ΔS of that bath. Note that if T was measured in energy units instead of an arbitrary temperature scale, there would be no need for a constant $k_B$, and entropy change could then be dimensionless, more in keeping with its basic relation to the change in the number of molecular states. Historically, though, as a physical property, temperature and ways to measure it preceded ideas about entropy.

A convenient way of thinking about the magnitude of an arbitrary entropy change ΔS is by asking: how much energy would have to be dissipated in a thermal bath with temperature T to produce the same entropy change ΔS? From Eq [A13], the dissipation of an energy ΔE equal to $k_B T$ produces an entropy change $\Delta S = k_B$. We can thus describe any entropy change in terms of the number of energy units $k_B T$ that would need to be dissipated to produce an equal entropy change.

Thinking of an entropy change $\Delta S$ in terms of the equivalent number of energy units $k_BT$ dissipated also provides a way to think about when an entropy change $\Delta S$ will have a significant effect on reducing the rate of a process through Eq [A11]: dissipation of one energy unit $k_BT$ creates an entropy change of one $k_B$, so the argument of the exponential in Eq [A11] is simply the number of dissipated energy units $k_BT$ that would produce an equivalent $\Delta S$. For example, if $\Delta S$ is equivalent to the dissipation of one energy unit $k_BT$, the entropy term in parentheses in Eq [A11] is (1-1/e) = 0.63. If $\Delta S$ is equivalent to dissipation of 3 $k_BT$ energy units, the net rate is reduced to $0.95R_0$, and if $\Delta S$ is equivalent to 5 $k_BT$ units the net rate is $0.99R_0$.

**Work.** The concept of 'work' essentially refers to a concentration of energy on a macroscopic scale, such as lifting a weight or turning a wheel. A simple way to think about this is to picture 'doing work' as just the transfer of energy to a system with no change in entropy of that system. For example, lifting a weight adds energy to the weight but does not change its entropy (the number of molecular states does not change if all of the molecules are lifted up). Taking the example above of an energy transfer $\Delta E$ between a system with temperature $T_1$ and a colder system with temperature $T_2$, there is a net increase in entropy. Suppose that this transfer of energy is now coupled to a process that does work (e.g., lifting a weight) involving a transfer of energy $E_W$ to a third system (the weight), with no change in entropy of that third system. Then less energy is transferred to the lower temperature bath. The net entropy change, coming from the energy transfers of the first and second thermal baths, is then:

$$\Delta S = -\frac{\Delta E}{T_1} + \frac{\Delta E - E_W}{T_2}$$

[A15]

As $E_W$ increases, $\Delta S$ is reduced, so the maximum extracted work will occur when $\Delta S$=0. The maximum fraction of the transferred energy $\Delta E$ that can be extracted as work is then:

$$\left(\frac{E_W}{\Delta E}\right)_{MAX} = 1 - \frac{T_2}{T_1}$$

[A16]

Operating an engine based on energy transfer from a hotter to a colder system works with better efficiency as the temperature difference increases. Note, though, that by the earlier arguments the net rate of the process would go to zero as ΔS goes to zero, so the maximum efficiency can only be attained as the rate goes to zero.

**Free energy**. Extracting work from the transfer of energy between two thermal baths is just one example of a general principle. If a process on its own has an entropy change ΔS, and if this process is then coupled to transfer of energy from a thermal bath at temperature T (involving a negative entropy change $-E_W/T$) to a macroscopic system (doing work—no entropy change), the maximum work $E_W$ that can be done corresponds to a net entropy change of zero. The net entropy change is $\Delta S - E_W/T$, so the maximum work is $E_W = T\Delta S$. For a chemical transformation we can use the general expression for the entropy change (Eq [A3]):

$$E_W = k_B T \ln \frac{\Phi}{\Phi_0}$$

[A17]

The maximum energy $E_W$ that can be extracted as work from a chemical process operating at a temperature T has historically been called the 'free energy'. This terminology can be confusing, though, as it suggests that there might also be 'bound energy', and there is no physical basis for this idea. The term 'free energy' likely originated with the general challenge of trying to understand why only some of the energy of a thermal bath can be extracted as work, even though from the point of view of energy conservation there would seem to be no limit. One could perhaps think of "free energy" as the amount of energy that could be "freed" from a thermal bath when coupled to another transformation with a positive entropy change. The essential idea that explains this, though, is entropy rather than energy, and from Eq [A17] free energy is a re-scaling of the entropy change. For this reason the free energy change, even though it has units of energy, does not behave like energy—it is not conserved—because it is essentially representing an entropy change. Note that free energy as the amount of work that could be extracted is equivalent to the earlier comment that we can think of a given entropy change $\Delta S$ in terms of the number of units of energy $k_B T$ dissipated in a thermal bath with temperature T that would create an equal entropy change $\Delta S$.

Two forms of free energy are usually distinguished, differing in the conditions under which the chemical transformation takes place, and so differing in the value $\Phi_0$. If volume is held constant, it is called the Helmholtz free energy, and if pressure is constant it is called the Gibbs free energy. In modern usage, to avoid the confusion of which free energy one is talking about, 'free energy' usually means the Helmholtz free energy, and 'Gibbs potential' is used for the Gibbs free energy.

Finally, the form of Eq [A17] is not the usual way that the Gibbs potential is expressed. First, it usually involves the gas constant R, rather than Boltzmann's constant $k_B$, to express the free energy in units of energy per mole of molecules undergoing the reaction. In addition, the ratios $\Phi$ and $\Phi_0$ are referred to a standard set of conditions defined by a ratio $\Phi_S$. The expression for the Gibbs potential $\Delta G$ is then:

$$\Delta G = RT \ln \frac{\Phi}{\Phi_S} - RT \ln \frac{\Phi_0}{\Phi_S}$$

[A18]

If the 'standard conditions' are defined such that $\Phi_S = 1$, the Gibbs free energy change can be written as:

$$\Delta G = \Delta G_0 + RT \ln \Phi$$

[A19]

where $\Delta G_0$ is the free energy change under the standard conditions. Although this formulation is common, as a mathematical expression it is problematic because in general $\Phi$ is not a dimensionless quantity: if the number of reactants and products is different, $\Phi$ will have dimensions related to concentration units that do not cancel. This means that the numerical value of the logarithm will depend on the specific units used to measure concentration. In contrast, the physics derivations used here lead to Eq [A3], where the ratios $\Phi$ and $\Phi_0$ have the same physical units, so the argument of the logarithm function is dimensionless regardless of the units used to measure concentration. To apply the usual form of the Gibbs potential, Eq [A19], one must be careful to use the same physical units for concentration that were used to make the ratio for the standard conditions $\Phi_S$ have a numerical value of one. That is, if the standard conditions are that each of the reactants and products has a concentration of 1 mole/liter, which will always make $\Phi_S$ have a numerical value of one, then for the calculation of $\Phi$ in Eq [A19] one must also measure all concentrations in moles/liter (e.g., using mM would lead to an inconsistent value of $\Phi$).

In principle, the basic idea of preserving the entropy change could be recast as preserving the free energy change, as this author described it in an early version of the idea [4]. The goal of the current work is to move past the more constricted idea of free energy (and whether it should be Gibbs or Helmholtz free energy) and the peculiar form in which it usually appears in chemical thermodynamics applications, and focus instead on the more fundamental notion of the entropy change.
