## Supplementary material for "The thermodynamics of thinking: connections between neural activity, energy metabolism and blood flow": Oxygen Transport Model

Submitted Nov 1, 2019

Paper for theme issue of *Philosophical Transactions of the Royal Society*, Part B:

'Significant relationships between non-invasive functional neuroimaging and the underlying neuronal activity'

Editors: Clare Howarth, Ralph Freeman, and Anusha Mishra

Corresponding author:
R.B. Buxton
University of California, San Diego, 9500 Gilman Drive, MC 0677, La Jolla, CA 92093-0677, USA

Funding sources:
This work was supported by the National Institutes of Health grants NS036722, MH111359, MH112969, and MH113295.

<u>Outline of the Supplementary Information</u>

**B. Model for the transport of $O_2$ and $CO_2$ to determine the mitochondrial ratio of $[O_2]/[CO_2]$**

B1. Parameters and units

B2. Overview of the model

B3. A more detailed discussion of the origin of Eq [B2]: Krogh cylinder model and the time constant $\tau$

B4. Blood Oxygen Content

B5. Transport of $CO_2$ and tissue $pCO_2$

B6. Estimating the Average Capillary $pO_2$

B7. Application of the model

**Additional notes for the calculations for the Figures in the main text**

**References**

### B. Model for the transport of $O_2$ and $CO_2$ to determine the mitochondrial ratio of $[O_2]/[CO_2]$

#### B1. Parameters and units

The key parameters, with units and typical values, are as follows. Blood flow (CBF) is abbreviated as $F$, and defined as the volume of arterial blood delivered to the capillary bed per minute divided by the total volume of the tissue element, and it is useful to think of $F$ as essentially having units of 1/min, with a typical human CBF of 50 ml blood/100g-min being about $F$ = 0.5 min$^{-1}$. The $O_2$ metabolic rate ($CMRO_2$) is abbreviated as $R_{O2}$, and is usually expressed as millimoles/100g-min. Here again the units can be simplified by noting that $CMRO_2$ essentially has dimensions of concentration/time (mM/min), with a typical value for human brain of about 1.8 mM/min. The oxygen extraction fraction (OEF) is dimensionless and abbreviated as $E$, and in the human brain a typical value is $E$=0.4. The hemoglobin concentration $H$ of arterial blood is expressed in milliequivalents of $O_2$ (i.e., the total concentration of $O_2$ that would result if the hemoglobin is fully saturated), and in these units a typical value is about 9 mM (e.g., in normoxia, when hemogobin bound $O_2$ is much larger than $O_2$ as dissolved gas, the total concentration $C_a$ of $O_2$ in arterial blood is about 9 mM; and H(mM) is equivalent to 0.62H( g/dL). Oxygen concentration of dissolved gas is expressed as an $O_2$ partial pressure ($PO_2$), in units of Torr (mmHg), with 7.5 Torr = 1 kPa. The solubility of $O_2$ in plasma is $\alpha_{O2}$ = 0.00134 mM $O_2$/Torr (equivalent to 0.003 ml $O_2$/dL-mmHg; i.e, 1 mM $O_2$ concentration = 2.24 ml $O_2$/100 ml). The $CO_2$ concentration as a dissolved gas is also expressed as a partial pressure, with a solubility $\alpha_{CO2}$ = 0.0307 mM/Torr.

#### B2. Overview of the model

The basic equation of mass balance for oxygen metabolism is:

$$R_{O2} = EFC_a \tag{B1}$$

The quantity $FC_a$ is simply the rate of $O_2$ delivery to a tissue element, and $E$ is then the fraction that is metabolized. Consistent with this relationship, we take as baseline normal values: $E = 0.4$, $F = 0.5$ min$^{-1}$, $C_a = 9$ mM, and $R_{O2} = 1.8$ mM/min.

Oxygen metabolism also can be considered as a diffusive flux of $O_2$ (as a dissolved gas) down a gradient from the $PO_2$ in the blood plasma to the $PO_2$ in the mitochondria, modeled as:

$$R_{O2} = \frac{\alpha_{O2}}{\tau}\left[P_C - P_T\right] \qquad \text{[B2]}$$

In this equation, $P_C$ is the average $PO_2$ in the capillary, $P_T$ is the average $PO_2$ in the tissue space, and $\tau$ is a time constant that depends on the diffusion coefficient of $O_2$ and the capillary bed geometry. Note that although this equation deals only with average values of $PO_2$, its derivation does not require that $PO_2$ is uniform in either the capillary or the tissue space. Section **B3** essentially serves as an appendix to the current section, providing a more detailed discussion of Eq [B2] with a derivation from the classic Krogh cylinder model with uniform tissue oxygen metabolism, which explicitly models spatial variability of $PO_2$. The simplicity of Eq [2] suggests that this basic relationship may apply for capillary beds with a more complicated geometry than the simple Krogh cylinder model, but this remains to be tested with more detailed computational modeling of realistic vascular beds. For now we take Eq [2] as an approximation describing the diffusive transport of $O_2$ from capillaries to tissue.

The parameter $\tau$ is characteristic of the capillary geometry and $O_2$ diffusivity, and captures the microscopic details of the diffusion process. A useful way to think about $\tau$ is that it is the parameter that embodies the effects of capillary density on $O_2$ transport: as capillary density increases, $\tau$ is reduced. The microscopic details that go into the determination of $\tau$ are not well known for the human brain, and are likely to vary across subjects and conditions. For this reason, in the calculations to follow we do not assume an *a priori* estimate of $\tau$, but rather treat $\tau$ as a parameter that is determined from the assumed baseline state by requiring consistency with Eqs [B1-2]. For simulating acute changes (e.g., neural activation) this value of $\tau$ is then assumed to remain constant (i.e., no acute capillary recruitment). For chronic changes, such as acclimatization to high altitude, the parameter $\tau$ may change.

In addition to the two equations above, the final key assumption of the model is that the tissue $PCO_2$ is equal to the venous $PCO_2$. A similar equilibration is not assumed for $O_2$ transport, and here the $CO_2$ equilibration is justified due to the much larger solubility of $CO_2$ in water than $O_2$, leading to larger fluxes of $CO_2$ with weaker gradients. The full transport model for $O_2$ and $CO_2$ is described in sections **B3-B6**, and the application of the model to calculate the tissue $[O_2]/[CO_2]$ ratio under different physiologic conditions is described in section **B7**.

### B3. A more detailed discussion of the origin of Eq [B2]: Krogh cylinder model and the time constant $\tau$

The Krogh cylinder model is a classical approach for modeling the exchange of $O_2$ with tissue. The capillary is treated as a long straight cylinder of radius $a$ and length $L$, with $O_2$ diffusing out into a larger concentric cylinder with radius $R$. Spatial coordinates are specified in terms of distance along the cylinder $z$ (ranging from 0 to $L$) and a radial distance $r$ (ranging from $a$ to $R$ for the extravascular space). The $O_2$ concentration in capillary plasma is expressed as an oxygen tension $pO_2$ (in mmHg), with a solubility $\alpha_{O2}$ that converts $O_2$ tension to $O_2$ concentration (mM).

The standard analysis focuses on a thin disk at $z$ in which $O_2$ is diffusing from a partial pressure $p_c(z)$ in the capillary, and the rate of $O_2$ consumption in a thin disk at $z$ is $M(z)$, and is assumed to be uniform within the disk. Radial diffusion of $O_2$ is assumed to be described by a diffusion coefficient $D$, and diffusion down the length of the cylinder is neglected. Solution of the diffusion equation for this scenario then provides an expression for the tissue oxygen tension $p_T(z,r)$ as a function of $r$ at different positions $z$ along the cylinder [1]:

$$p_T(z,r) = p_c(z) - \frac{M(z)}{2s_{O2}D}\left[R^2 \ln\frac{r}{a} - \frac{(r^2 - a^2)}{2}\right] \qquad \text{[B3]}$$

Note that this analysis does not specify how the capillary $O_2$ partial pressure $pO_2$ varies along the capillary, but only how tissue $pO_2$ varies with radial distance for a given capillary $pO_2$ profile $p_c(z)$. Based on this formulation, we can define an average extravascular $O_2$ tension $p(z)$ by averaging Eq [B3] over $r$, and this can be written as:

$$p(z) = p_c(z) - \frac{m(z)}{s_{O2}}\tau \qquad \text{[B4]}$$

where $\tau$ is a time constant defined by the cylinder geometry and the diffusion coefficient:

$$\tau = \left\langle \frac{R^2 \ln\frac{r}{a} - \frac{(r^2 - a^2)}{2}}{2D} \right\rangle \qquad \text{[B5]}$$

where the bracket indicates the average over the range $r = a$ to $r = R$. To give a sense of the magnitude of $\tau$, the average in Eq [B5] can be calculated numerically. For a capillary radius $a$=3μ, a cylinder radius $R$=30μ, and a diffusion coefficient $D$=1 $m^2$/ms, $\tau$ is 0.71 seconds.

The variation of capillary $Po_2$ along the capillary is determined by mass balance. Let $C(z)$ be the total blood concentration of $O_2$ at position $z$, and let $u$ be the speed of blood moving down the capillary (assumed uniform in the capillary). In a time interval *dt*, the blood moves a distance *dz=udt*, so we can consider a disk with thickness *dz*. For a steady-state, during *dt* the amount of $O_2$ consumed in the disk must match the amount of $O_2$ lost from the blood:

$$\mathrm{O_2\,consumed} = M(z)\,dt\,dz\,\pi(R^2 - a^2) \qquad [B6]$$

$$\mathrm{O_2\,lost} = \left(-\frac{dC}{dz}dz\right)\pi\, a^2\, dz \qquad [B7]$$

So the basic equation for total capillary $O_2$ is:

$$\frac{dC}{dz} = -\frac{R^2 - a^2}{a^2 u} M(z) \qquad [B8]$$

The cerebral blood flow *F* (ml blood/s-ml tissue) is the rate of delivery of blood to the cylinder, divided by the cylinder volume:

$$F = \frac{a^2 u}{R^2 L} \qquad [B9]$$

At this point it is convenient to express the distance along the capillary as a normalized variable $x=z/L$, with *x* ranging from 0 to 1. If we further assume that $R^2 >> a^2$, then Eq [B8] can be written as:

$$\frac{dC}{dx} = -\frac{M(x)}{F} \qquad [B10]$$

Based on the form of Eq [B2] suggested by the Krogh model, we have:

$$M(x) = \frac{s_{O2}\left[p_c(x) - p(x)\right]}{\tau} \qquad [B11]$$

Combining Eqs [B10] and [B11] to eliminate *M*(*x*) gives:

$$\frac{dC(x)}{dx} = -\frac{\alpha_{O2}}{F\tau}\left[p_c(x) - p(x)\right] \quad \text{[B12]}$$

The term *M*(*x*) was eliminated between these equations, but note that this does not assume that *M*(*x*) is uniform along the cylinder in this analysis.

Finally, Eq [B2] results from averaging Eq [B12] over *x*. The average of $p_c(x)$ is just the average capillary $PO_2$, and the average of p(x) is the average $PO_2$ within the tissue cylinder, and the average value of the derivative of the blood concentration is –ECa, giving:

$$EC_a = \frac{\alpha_{O2}}{F\tau}\left[P_C - P_T\right] \quad \text{[B13]}$$

**B4. Blood Oxygen Content**

In blood, the $O_2$ concentration in the form of dissolved gas is a small fraction of the total $O_2$ concentration because the majority of $O_2$ is bound to hemoglobin. The hemoglobin concentration in arterial blood *H* is expressed in equivalents of $O_2$ for full saturation (mM), and the fractional $O_2$ saturation of hemoglobin $S_{O2}$ ranges from 0 to 1. As plasma $O_2$ diffuses out of the capillary, $O_2$ from the hemoglobin is assumed to quickly reach a new equilibrium with the plasma, so that the total $O_2$ concentration is:

$$C = \alpha_{O2}P_{O2} + S_{O2}H$$

[B14]

The hemoglobin saturation is assumed to be related to the blood plasma $pO_2$ by the Hill equation:

$$S_{O2} = \frac{1}{1 + \left( {}^{P_{50}}/_{P_{O2}} \right)^{h}}$$

[B15]

with $h$=2.8 and $p_{50}$=27 mmHg taken as typical values for human blood [2]. The equivalent expression for $PO_2$ is:

$$P_{O2} = P_{50} \left( \frac{S_{O2}}{1 - S_{O2}} \right)^{1/h}$$

[B16]

The value of $P_{50}$ defines the plasma $PO_2$ that produces 50% hemoglobin saturation, and this parameter is important because it determines the average capillary $PO_2$ for a given oxygen extraction fraction $E$—a higher $p_{50}$ for the same $E$ will correspond to a higher capillary $PO_2$. However, $P_{50}$ does not remain constant down the length of the capillary because of the Bohr effect: $P_{50}$ increases as $CO_2$ is taken into the capillary blood, reducing blood pH. This effect is incorporated in the model with the assumption that $P_{50}$ varies with pH as:

$$P_{50} = P_{50a}\, 10^{-b\,(\mathrm{pH}\text{-}pH_a)}$$

[B17]

where $P_{50a}$ is the value on the arterial end of the capillary and $pH_a$ is the arterial pH. The parameter $b$ describes the magnitude of the Bohr shift, and a typical value in human blood is 0.48 [3], which is assumed in all of the model calculations.

#### B5. Transport of $CO_2$ and tissue $pCO_2$

A model for the transport of $CO_2$ is needed in part because the goal is to model changes in tissue $PCO_2$, and also because additional $CO_2$ carried in blood will affect the pH that in turn affects $P_{50}$ and the average capillary $PO_2$. Carbon dioxide diffuses from tissue into blood as dissolved gas, but once in the blood the majority is converted to bicarbonate ions ($HCO_3^-$) in plasma, catalyzed by the enzyme carbonic anhydrase. In addition, $CO_2$ binds to deoxyhemoglobin to form carbamino hemoglobin, which is a significant carrier of $CO_2$ in venous blood. This complexity of $CO_2$ carriage in blood makes the relationship between total $CO_2$ ($T_{CO2}$) and $PCO_2$ somewhat complicated, depending on hemoglobin content (H), pH, and hemoglobin saturation $S_{O2}$. Douglas and colleagues developed the following empirical model for $T_{CO2}$ [4]:

$$T_{CO2} = C_{CO2}\left[1 - \frac{.0466\,H}{(3.352 - 0.456\,S_{O2})(8.142 - pH)}\right]$$

[B18]

where $C_{CO2}$, the 'total plasma $CO_2$', is given by the Henderson-Hasselbalch equation:

$$C_{CO2} = \alpha_{CO2} P_{CO2}(1 + 10^{pH-pK})$$

[B19]

with pK=6.091. Assuming that metabolized $O_2$ is all converted to $CO_2$, the arterial (a) and venous (v) values of total $CO_2$ are related by:

$$T_{CO2}(v) = T_{CO2}(a) + EC_a$$

[B20]

Finally, we need to consider the pH change between the arterial value and the venous value. Blood pH changes down the length of the capillary as $CO_2$ is added, equivalent to adding an acid, but the buffering in the blood also changes with $S_{O2}$ through the effects of hemoglobin. Rather than trying to model these separate effects, we used a simple empirical expression as a linear dependence on total $CO_2$ content of blood ($T_{CO2}$):

$$pH = pH_a - c\,[T_{CO2} - T_{CO2}(a)]$$

[B21]

In an early study with simultaneous measurements of blood gas values in an artery (femoral, radial or brachial) and internal jugular vein of resting human volunteers (male, mostly medical students), Gibbs et al [5] found a mean increase of total $CO_2$ from arterial to venous blood of 3.0 mM and a corresponding mean pH change of -0.053, giving an estimate for the value of *c* of about 0.018 for this approximation (for $T_{CO2}$ measured in mM).

In applying the above equations to model parameters for venous blood, the oxygen extraction fraction E and the arterial $O_2$ content $C_a$ determine the change in $T_{CO2}$ going from arterial to venous blood, which then determines the change in pH, and then these changes determine $P_VCO_2$. The tissue $PCO_2$ is assumed to be equal to $P_VCO_2$.

### B6. Estimating the Average Capillary $pO_2$

In order to apply Eq [2], the average capillary $pO_2$ ($P_C$) is needed. The key assumption is that the average capillary $pO_2$ is the $pO_2$ after half of the net extracted $O_2$ has been lost from the blood and half of the net acquired $CO_2$ has been added to the blood [2]. At this midpoint, the capillary $O_2$ concentration is:

$$C_c = \left(1 - \frac{E}{2}\right) C_a$$

[B22]

The capillary pH is estimated for the expression for pH in terms of added total $CO_2$ with Eq [B21], the parameter $P_{50}$ in the Hill equation is updated based on the pH, and the capillary $PO_2$ ($P_C$ in Eq [B13]) is then calculated.

### B7. Application of the model

The model is applied by starting with a defined baseline state with specified values of the following parameters: blood flow (F), oxygen metabolism ($R_{O2}$), hemoglobin content (H), arterial blood gases ($P_aO_2$ and $P_aCO_2$), arterial pH ($pH_a$), and tissue $PO_2$. **Table 1** lists the values for the standard baseline model used in this paper. Additional baseline parameters are calculated from the given parameters using the equations developed in the previous sections, including the baseline tissue ratio $[O_2]/[CO_2]$. These additional baseline parameters are listed in **Table 2**. A new state is then specified, altering some of the physiologic parameters (e.g., increasing $R_{O2}$), and the new tissue $[O_2]/[CO_2]$ is calculated. Then the value of blood flow (F) needed to maintain the original tissue $[O_2]/[CO_2]$ ratio in the new state is calculated.

In more detail, the added $CO_2$ to venous blood is assumed to be equal to the amount of metabolized $O_2$ (Eq [B20]). The $CO_2$ added to blood alters the blood pH (Eq [B21]). The $O_2$ binding to hemoglobin is modeled with the Hill equation, which is defined in terms of the $PO_2$ for which the saturation of hemoglobin is 50% (designated $P_{50}$), and the value of $P_{50}$ in turn depends on the blood pH (Eq [B17]). The mean blood $PO_2$ for use in Eq [2] is assumed to be the $PO_2$ of blood after half of the net total $CO_2$ generated by metabolism has been added (Eq [B22], i.e., approximating a capillary $PO_2$). Specifically, the capillary $PO_2$ is estimated by removing a total fraction E/2 of the $O_2$ from the arterial blood to estimate the total $O_2$ change, adding the same amount of $CO_2$ and calculating the associated blood pH change, adjusting the value of $p_{50}$ for the pH change, and calculating the corresponding $PO_2$. Then the tissue $O_2$ is determined from Eq [B2].

As a first step, this procedure is applied to the assumed baseline values, and because the baseline tissue $pO_2$ is assumed, these calculations will define the value of $\tau$ to be consistent with Eq [2], and for subsequent changes in state that value of $\tau$ is assumed to remain constant and the basic parameters are recalculated for the new state in a similar way. Note that the assumption of constant $\tau$ for the new state is essentially an assumption of unchanged diffusion properties and geometry of the capillary bed (no capillary recruitment [6]), but for modeling long-term hypoxia effects it should be considered that

$\tau$ could be shortened by growth of new blood vessels. To estimate tissue $PCO_2$, the venous $PCO_2$ ($P_VCO_2$) is calculated with Eq [B18] relating total $CO_2$, $SO_2$, pH and $P_VCO_2$.

**Table 1**. Reference baseline state for the human brain: assumed parameter values

| **Parameter** | **Definition** | **Assumed Baseline Value** |
|---|---|---|
| $F$ | Cerebral blood flow | 0.5 $min^{-1}$ (45 mL/100mL-min) |
| $E$ | $O_2$ extraction fraction | 0.4 |
| $P_T$ | Tissue $O_2$ partial pressure | 25 Torr |
| $H$ | Arterial hemoglobin | 9 mM $O_2$ at 100% saturation |
| $pH_a$ | Arterial pH | 7.4 |
| $PaO_2$ | Arterial $O_2$ partial pressure | 100 Torr |
| $PaCO_2$ | Arterial $CO_2$ partial pressure | 40 Torr |
| $\alpha_{O2}$ | $O_2$ solubility | 0.00135 mM/Torr |
| $\alpha_{CO2}$ | $CO_2$ solubility | 0.0307 mM/Torr |
| pK | Effective pK for CO2/bicarbonate | 6.1 |
| $h$ | Hill equation exponent | 2.8 |
| $P_{50a}$ | arterial half-saturation of hemoglobin | 27 Torr |
| $b$ | Bohr shift parameter | 0.48 |
| $c$ | parameter for pH dependence on $T_{CO2}$ | 0.018 $mM^{-1}$ |

**Table 2**. Reference baseline state for the human brain: calculated parameter values from the model for the baseline state defined by Table 1

| **Parameter** | **Definition** | **Typical Value** |
|---|---|---|
| $R_{O2}$ | Cerebral metabolic rate of $O_2$ | 1.78 mM/min (4 mL $O_2$/100mL-min) |
| $C_a$ | Arterial blood $O_2$ concentration | 8.9 mM |
| $T_{CO2}$(a) | Total arterial total $CO_2$ | 20.7 mM |
| $pH_v$ | Venous pH | 7.34 |
| $P_C$ | Average capillary $O_2$ partial pressure | 44.5 Torr |
| $P_TCO_2$ | Average tissue $CO_2$ partial pressure | 52.3 Torr |
| $\Phi_{O2base}$ | Baseline tissue ratio $[O_2]/[CO_2]$ | 0.021 |
| $\tau$ | $O_2$ diffusion time constant | 0.0147 min |

**Additional notes for the calculations for the Figures in the main text**

**Figure 3A**: In the study of Nioka et al [7], the 'baseline' normoxic state was actually more of a hyperoxic state (arterial $PO_2$ was ~145 mmHg). In the model calculations the same normal baseline state was used as in the other calculations (arterial $PO_2$ = 100 mmHg). In addition, no modification of the P50 of $O_2$ binding to hemoglobin was made, which may be slightly different in dogs compared to humans. In the model calculations the stages of hypoxia were taken to be defined by the reported arterial $PO_2$ values, rather than the reported $O_2$-hemoglobin saturation values, which is why the model points have slight difference in $O_2$ saturation from the measured data points. Note also that the current interpretation of the data here in terms of the transport model differs from the interpretation of the original authors, who assumed tissue $O_2$ concentrations were much lower than what the transport model indicates.

**Figure 4B:** For comparison between the measured alveolar values and the arterial values used in the model, the arterial and alveolar $PCO_2$ values were assumed to be the same. The alveolar $PO_2$ value was assumed to be 3 mmHg higher than the arterial value, based on the studies by Crapo and colleagues [8]. The shift is likely to increase slightly to about 5 mmHg as altitude increases: Crapo and colleagues found an increase to 5 mmHg at 1400m elevation, and Grocott and colleagues [9] found a shift of 5.4 mmHg for subjects on Mount Everest. For a more complete analysis of these high altitude acclimatization effects this aspect should be modeled as well.
