## Supplementary material for "The thermodynamics of thinking: connections between neural activity, energy metabolism and blood flow": Neural Model

Submitted Nov 1, 2019

Paper for theme issue of *Philosophical Transactions of the Royal Society*, Part B:

'Significant relationships between non-invasive functional neuroimaging and the underlying neuronal activity'

Editors: Clare Howarth, Ralph Freeman, and Anusha Mishra

Corresponding author:
R.B. Buxton
University of California, San Diego, 9500 Gilman Drive, MC 0677, La Jolla, CA 92093-0677, USA

Funding sources:
This work was supported by the National Institutes of Health grants NS036722, MH111359, MH112969, and MH113295.

### Outline of the Supplementary Information

**C. Neural model connecting excitatory and inhibitory activity to $CMRO_2$ and CBF**

**Additional notes for the calculations for the Figures in the main text**

**References**

### C. Neural model connecting excitatory and inhibitory activity to $CMRO_2$ and CBF

To begin to model how different aspects of neural activity could drive $CMRO_2$ and CBF, we used a variant of the Wilson-Cowan (WC) model [1] of interacting populations of excitatory (E) and inhibitory (I) neurons to explore the increase in E and I activity as input to the E population increases. We used a simplified form of the WC model in which the dynamic behavior is described by:

$$\tau_0 \frac{dE}{dt} = -E + S(wE - wI + P)$$

$$\tau_0 \frac{dI}{dt} = -I + S(wE + Q)$$

*S*(*x*) is a sigmoid function, taken to be:

$$S(x) = \frac{1 - e^{-ax}}{1 + e^{a(x-\theta)}}$$

when $x>0$ and $S(x)=0$ when $x<0$. The parameters used were $a=2$, $\theta=2$ and $\tau_0=10$ms. The parameter *P* is the input to the E population, and the parameter *Q* is the input to the I population. The weight *w* connecting the different components was for simplicity taken to be the same, but in principle each instance of *w* in the equations above could take on a different value. Note that "$\tau_0$" in this section has nothing to do with the "$\tau$" in **Section B**.

#### Additional notes for the calculations for the Figures in the main text

**Figure 5B:** For the Wilson-Cowan model calculations, the input to the excitatory population (E) was varied as shown on the x-axis (corresponding to the parameter P in the model). The input to the inhibitory population (I) was held constant at a value of 0.2 (the parameter Q in the model). The differential equations of the model were integrated for 4000 time steps of 1ms to allow the populations to settle to a steady-state. Note that a self-inhibitory signal from the I population back to itself was not included, although such a connection was included in the original WC model formulation.

**Figure 5D:** The BOLD curves for the two models were calculated in a way consistent with the calibrated BOLD estimates of $CMRO_2$ in the original paper [2], using the Davis model with $\alpha$=0.4 and $\beta$=1.5. The BOLD curve calculated for the WC model is actually slightly negative for the lowest CBF change, but by a negligibly small amount (~0.01%).
